# Identifying Modulators of Cellular Responses by Heterogeneity-sequencing

**DOI:** 10.1101/2024.10.28.620481

**Authors:** Kevin Berg, Lygeri Sakellaridi, Teresa Rummel, Thomas Hennig, Adam W Whisnant, Manivel Lodha, Tobias Krammer, Christophe Toussaint, Katarzyna Szymanska-De Wijs, Yan Zheng, Bhupesh K Prusty, Lars Dölken, Antoine-Emmanuel Saliba, Florian Erhard

## Abstract

The response of individual cells to drug treatment, virus infections or other molecular stimuli is highly heterogeneous and depends on the cell’s initial state. Library preparation for single-cell transcriptomics is destructive, precluding a direct comparison between the initial state and the stimulus outcome. Consequently, current methods are restricted to identifying correlative associations rather than resolving causal drivers of heterogeneous outcomes. We developed Heterogeneity-seq, which combines single-cell RNA-seq with metabolic RNA labeling (scSLAM-seq) and double machine learning to overcome this limitation. By leveraging simultaneous measurements of unlabeled and labeled RNA in individual cells, Heterogeneity-seq uncovers the transition from pre-stimulated cell states to distinct stimulation outcomes across thousands of cells. These links enable the identification of factors that causally govern heterogeneous cellular responses. We used Heterogeneity-seq to identify both known and novel genes that drive responses to drug treatment, as well as pro- and antiviral host factors governing cytomegalovirus infection.

## Introduction

Genetically identical cells can exhibit remarkably diverse responses to external challenges such as drug treatment or virus infection ^1,2^. These divergent outcomes depend on distinct cellular states that also exist in a seemingly homogeneous population of cells. Finding causal factors in these cell states that determine outcomes is crucial for deciphering the molecular basis of cellular decision-making and predicting cellular responses to external stimuli.

So far, molecular factors governing such responses have been identified using active perturbation strategies, such as RNA interference via short hairpin RNA or genome editing using CRISPR/Cas9 technology ^3,4^. These approaches systematically disrupt candidate genes to assess their impact on cellular phenotypes. While such methods yield valuable insights into gene function and cellular response pathways, they are not without limitations. Active perturbations introduce artificial changes that may not fully recapitulate physiological conditions. For instance, RNAi-mediated knockdown can induce off-target effects, activate cellular stress responses, or trigger compensatory pathways that obscure the direct effects of gene silencing ^5^. Similarly, CRISPR-based genome editing, while precise, can lead to unintended genomic alterations or adaptation mechanisms that complicate the interpretation of results ^6^. Moreover, for all these approaches the active perturbation of candidate genes is performed days before analyzing the stimulus and cellular response and, thus, cells may undergo extensive changes in state, potentially masking the true role of the targeted factor. As a result, the observed effects might stem from indirect regulatory influences of the perturbation rather than direct causation.

Within any population of genetically identical cells, gene expression is inherently heterogeneous — that is, the abundance of individual transcripts and proteins fluctuates from cell to cell, even within the same cell type and under uniform environmental conditions. This heterogeneity arises from the stochastic nature of transcription and translation, differences in cell-cycle stage, and subtle microenvironmental influences ^7,8^. As a result, what appears to be a homogeneous culture indeed contains a spectrum of transcriptional and phenotypic states, which contributes to the functional properties of the respective tissue ^9^. If both the causal factor and the molecular outcome of the external stimulus can be monitored for each individual cell over time from before the stimulus up to a time where the outcome is observable for individual cells, active perturbation of the factor is not required. Instead, expression heterogeneity can be considered a combined loss- and gain-of-function screen at physiological levels: Cells that show weak or strong expression of a causal factor before stimulation will differ in their response to the challenge. Thus, positive or negative correlation between gene expression before challenge to the molecular outcome of the challenge predicts a positive or negative impact of the gene.

This concept is fundamental to clone tracing methods, that have been used to retrospectively identify cells that are susceptible to drug treatment or virus infection ^10,11^. Clone tracing is based on introducing DNA barcodes into cells, expanding them and applying the stimulus to a subset of the population. The barcodes are then used to link a stimulated cell having a specific outcome to a cell before stimulus representing the prior state. Such approaches successfully identified states governing outcomes that are preserved in cell clones over multiple cell divisions ^10,11^. However, clone tracing is blind towards more transient states, including the cell cycle or stochastic resistance to BRAF inhibitor treatment in melanoma ^12^ or the fluctuating abundance of sensors and signal transducers in the interferon response ^13,14^.

A major limitation of scRNA-seq is that each single cell can only be analyzed once, either before or after a stimulus. However, metabolic RNA labeling combined with chemical nucleotide conversion sequencing and scRNA-seq ^9,15,16^ provides an elegant means to circumvent this limitation: If labeling is initiated prior to the stimulus, unlabeled RNA reflects the prior state, whereas labeled RNA corresponds to the outcome. We call this concept of using metabolic labeling information to connect heterogeneous pre-stimulation states to heterogeneous stimulation outcomes “Heterogeneity sequencing” (Heterogeneity-seq).

Here, we demonstrate that Heterogeneity-seq can be implemented using scSLAM-seq or sci-fate ^9,15^. We present an optimized experimental protocol for scSLAM-seq on the 10x Chromium platform providing >35,000 UMIs per cell. We introduce our improved computational methodology GRAND-SLAM 3.0 ^17^, allowing us to reliably and efficiently quantify labeled and unlabeled RNA and associated uncertainties for a large-scale run. We developed standard analysis workflows for analyzing single-cell data, including labeled and unlabeled RNA by interfacing our grandR package ^18^ with Seurat ^19^. Heterogeneity-seq and all computational methods required for it are implemented in the publicly available HetSeq package. Using all these computational approaches, we show that Heterogeneity-seq can find known and novel factors that causally impact drug responses and infection outcomes using publicly available sci-fate data and newly obtained scSLAM-seq data, respectively.

## Results

### Heterogeneity-seq overview

To develop the Heterogeneity-seq approach, we first focused on published sci-fate data consisting of overall n=6,680 A549 cells that were treated with dexamethasone (DEX) to stimulate the glucocorticoid receptor ^15^. The experiment consisted of consecutive snapshots obtained at 2/4/6/8/10h after treatment, each labeled for 2h using 4-thiouridine (4sU). In addition, an untreated control with 2h labeling was performed (0h time point). After applying our GRAND-SLAM pipeline ^17,18^, we obtained a median of n=27,414 UMIs per cell.

At the total RNA level, cells from the 0h and 2h time points did not segregate in a UMAP (**Fig. 1a**), but were clearly separated at later time points, indicating that DEX treatment induces significant changes in the transcriptome. To quantitatively assess the strength of the DEX response, we computed a DEX score per cell reflecting the overall strength of induction of a defined set of n=105 DEX responder genes (**Fig. 1b**). These genes are targets in A549 cells of a module of 10 transcription factors including *CEBPB*, *FOXO1* and JUNB that have been described as downstream effectors of the glucocorticoid response ^20–22^. At the total RNA level, the DEX score gradually increased over the whole period (0-10h) (**Fig. 1c**). This gradual increase could be due to (***i***) a rapid response including a simultaneous activation of transcription of the responder genes in all cells within the first 2h followed by a slow increase of total RNA due to slow RNA turn-over kinetics, or (***ii***) an asynchronous and gradually increasing response of individual cells. Analyzing the DEX score at the level of labeled, newly synthesized (new) RNA revealed that the overall response occurred immediately during the first 2h. However, also in between 2h and 4h, but not later, there was a further induction of response gene expression (**Fig. 1c**). The extent of response gene expression was highly heterogeneous with weak responder cells (the bottom third with respect to the DEX score) having 2-fold less induction of average response gene expression than strong responder cells (the top third) at 2h and later (**Fig. 1d**).

**Figure 1.**
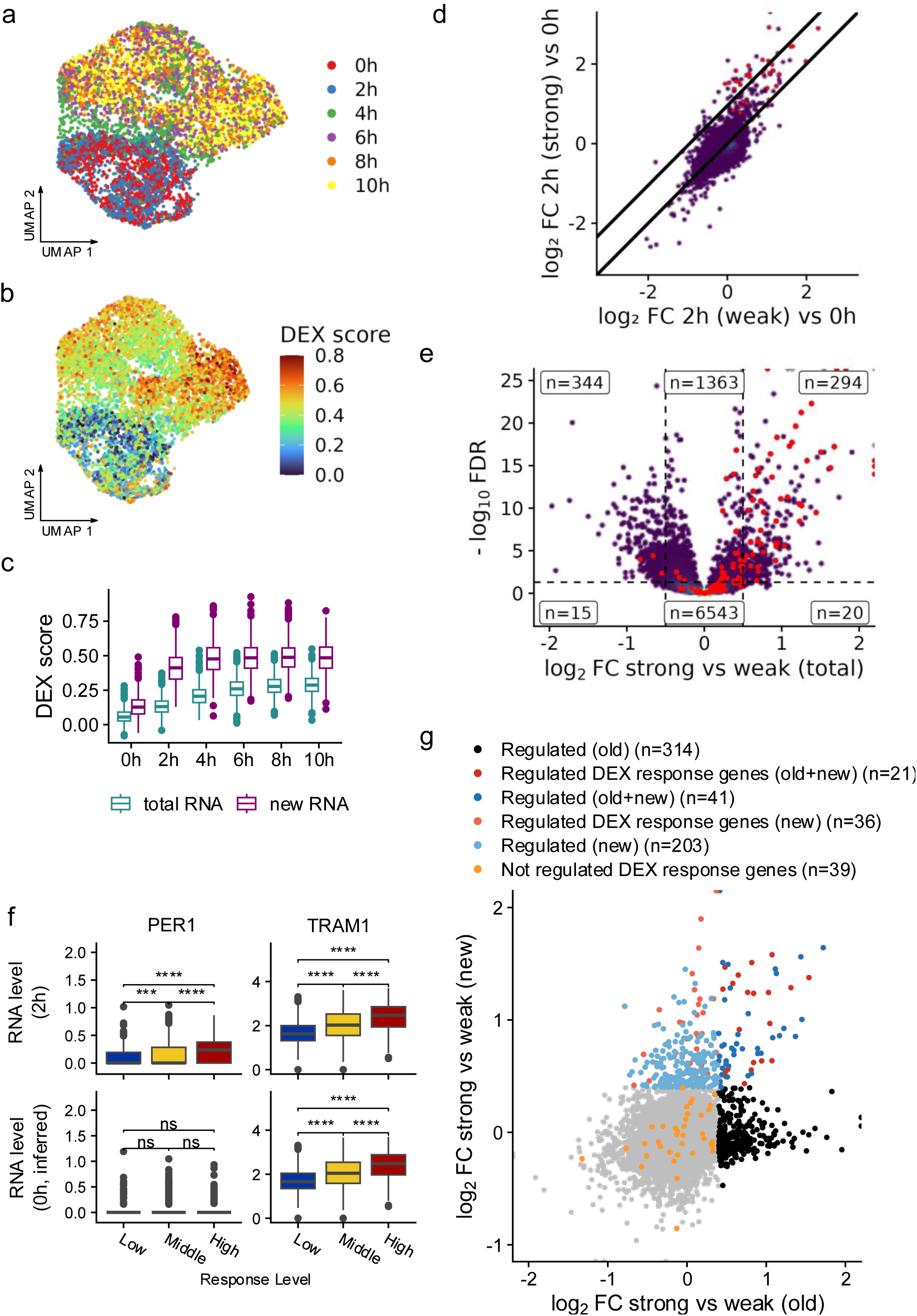
Gene regulation upon DEX treatment. **a** UMAP embedding of n=6,680 DEX-treated A549 cells from different time points as indicated **b** Same UMAP embedding as in a color according to DEX score. A higher DEX score indicates strong expression at total RNA level of DEX-inducible genes. **c** Boxplots (center line, median; box limits, upper and lower quartiles; whiskers, 1.5x interquartile range; points, outliers) showing DEX score distribution of all cells per time point. The DEX score was computed on both total and new RNA level as indicated. **d** Scatterplot of n=16,046 genes comparing log_2_ fold changes (FC) of a pseudobulk of weak responder cells (bottom third DEX scores at 2h) versus a pseudobulk of all 0h cells to log_2_ fold changes of a pseudobulk of strong responder cells (top third DEX scores at 2h) versus a pseudobulk of 0h cells (new RNA level). DEX response genes that were used to compute the DEX score are highlighted in red. The two lines show the line through origin indicating no differential expression, and the linear regression line of the DEX response genes. **e** Vulcano plot showing log_2_ fold changes of strong (top third DEX score) versus weak (bottom third DEX score) responder cells and corresponding p values (Wald test by DESeq2, Benjamini-Hochberg corrected). Thresholds on log_2_ fold changes (−0.5 and 0.5) and p values (0.05) as well as gene numbers belonging to corresponding groups are indicated. Genes that are induced at new RNA level at 2h (n=120, p<0.05, Wald test, Benjamini-Hochberg corrected, log_2_ FC > 0.5) are indicated in red. **f** Boxplots (center line, median; box limits, upper and lower quartiles; whiskers, 1.5x interquartile range; points, outliers) showing gene expression at total RNA level (log-normalized, Seurat, top row) of 2h cells stratified by DEX score (low, middle, high response level), or at half-life corrected old RNA level (bottom row) reflecting previous cell states for the genes PER1 and TRAM1. **g** Scatterplot comparing log_2_ fold changes of pseudobulks of 2h cells versus 0h cells at the old RNA level to the new RNA level. Genes are stratified as indicated by colors.

We first focused on the immediate response to identify factors associated with either strong or weak responses within the first 2h (subsequently called “HetSeq factors”). If a factor that positively impacts the response strength has stronger expression before treatment in one cell than in another, the former cell should respond more strongly to the treatment than the latter. Thus, an approach to find such positive or negative factors is to analyze gene expression before treatment in cell subsets that later show a strong or weak response.

Under the assumption that gene expression in these subsets does not change significantly at the total RNA level within 2h (**Fig. 1a**), we identified genes that are differentially regulated between cells with low (bottom third response score) or high (top third) response levels (**Fig. 1e**). Overall, there were n=344 genes that were stronger expressed in low response level cells and therefore candidates of negative factors, and n=294 genes that were stronger expressed in high response level cells representing candidates of positive factors (at least 1.41-fold change, P<0.05, two-sided Wilcoxon test, multiple testing corrected by Benjamini-Hochberg). Among these 294 genes, there was the known coregulator of the glucocorticoid response TRAM1 ^23^, but also response genes such as PER1 (**Fig. 1f, top**). Overall, among the genes differentially regulated between low and high responder cells there were n=54 out of 120 genes that were also regulated from 0h to 2h upon DEX treatment (at least 1.41-fold upregulated in new RNA, P<0.05, two-sided Wilcoxon test, multiple testing corrected by Benjamini-Hochberg, **Fig. 1e**). Taken together, this indicates that among the identified total RNA differential genes between low and high response level, there are indeed HetSeq factors, but also genes that are quickly regulated already within 2h.

To remove genes that are quickly regulated but not HetSeq factors, we performed differential gene expression analyses of unlabeled, pre-existing (old) RNA from the 2h cells reflecting the prior cell state before treatment. Old RNA was corrected for RNA degradation during the 2h of labeling (see Methods), and corrected old RNA is called ‘previous RNA’. Interestingly, for the response gene PER1, there was no difference in gene expression at the previous RNA level, whereas for the known positive factor TRAM1, the difference already observed at the total RNA level was also present in the previous RNA (**Fig. 1f, bottom**). Thus, for PER1, the observed differences at the total RNA level at 2h are a consequence and not the cause of a stronger response to DEX treatment. By contrast, since previous RNA levels reflect the prior cell states, this analysis suggests heterogeneous TRAM1 expression has an impact on the single cell level treatment outcome.

For a more comprehensive analysis, we decomposed changes observed at the total RNA level (**Fig. 1e**) into changes at the old RNA and at new RNA level, reflecting the prior state and response, respectively. In this analysis, response genes should show differential expression at the new RNA level, whereas HetSeq factors should have differential expression at the old RNA level. Focusing on upregulated genes, there were several groups (**Fig. 1g**): There were n=301 genes that showed induction of new RNA expression in high response level cells (at least 1.32-fold induced), which included n=57 of the predefined DEX response genes. Of the 301 genes, n=62 (21%) also showed differential expression at the old RNA level, which included n=21 DEX response genes. Furthermore, a separate group of genes exhibited only differential expression at the old RNA level (n=314). These 314 genes are prime candidates of positive HetSeq factors: For cells that later (at 2h) showed strong responses, before the treatment (old RNA), they were more strongly expressed than in cells that later showed weak responses. It remained, however, unclear from that analysis whether the 62 genes that showed differential expression in both new and old RNA were also HetSeq factors, or whether regulation in old RNA was only observed as spill-over from new RNA, since metabolic RNA labeling cannot always unambiguously differentiate between old and new RNA ^17^.

In conclusion, these analyses show that old RNA levels can, in principle, be used to identify HetSeq factors whose heterogeneous expression before drug treatment impacts the strength of the molecular response. However, spillover of new RNA into the old RNA compartment might occur due to technical limitations of metabolic RNA labeling in single cells, such that genes that are regulated might also appear as HetSeq factors.

### Convolution approach to NTR estimation

In nucleotide conversion RNA-seq experiments, such as sci-fate or scSLAM-seq ^9,15^, chemical conversion of 4sU residues incorporated during transcription induces T-to-C mismatches, which can be leveraged to quantify unlabeled and labeled RNA. However, typically only 2-8% of all U are substituted by a 4sU during transcription in the phase of labeling. Thus, counting reads with T-to-C mismatches underestimates labeled RNA ^24^. Additionally, substitutions due to sequencing errors or errors during reverse transcription or PCR can further introduce T-to-C mismatches for old RNA, resulting in overestimations of new RNA.

GRAND-SLAM is based on a mixture modeling approach. Instead of counting reads (or UMIs) with or without T-to-C mismatches to estimate new or old RNA expression, respectively, the model probabilistically assigns reads to old and new RNA, respecting both sequencing errors and other reasons that might induce T-to-C mismatches in old RNA, as well as the limited 4sU incorporation that results in reads without T-to-C mismatches from new RNA ^25^. The output of GRAND-SLAM is the new-to-total RNA ratio (NTR) for each gene, i.e., the percentage of labeled RNA among all RNA sequenced.

This model can provide unbiased and accurate estimates if there are reads from which information can be borrowed to recognize how many reads originate from a labeled RNA molecule but do not have T-to-C mismatches, which is typically the case for bulk RNA-seq (**Fig. 2a**). However, if reads originate from distinct cells, sharing information does occur to a lesser extent and labeled RNA might be underestimated if the mixture modeling approach is applied for individual cells (**Fig. 2b**). To test this, we estimated the overall NTR for all genes expressed in the 0h sample of the DEX treatment experiment by two means: First, we applied the mixture model for each cell individually and then pooled estimated new and old RNA for all cells. Second, we pooled all UMIs from all cells, and then applied the mixture model to estimate the pseudo-bulk NTR. This revealed that the two estimates diverged substantially for the majority of genes (11,282 out of 16,044 genes in the dataset, 70.3%) that were expressed on average with less than 1 copy detected per cell (**Fig. 2c**).

**Figure 2:**
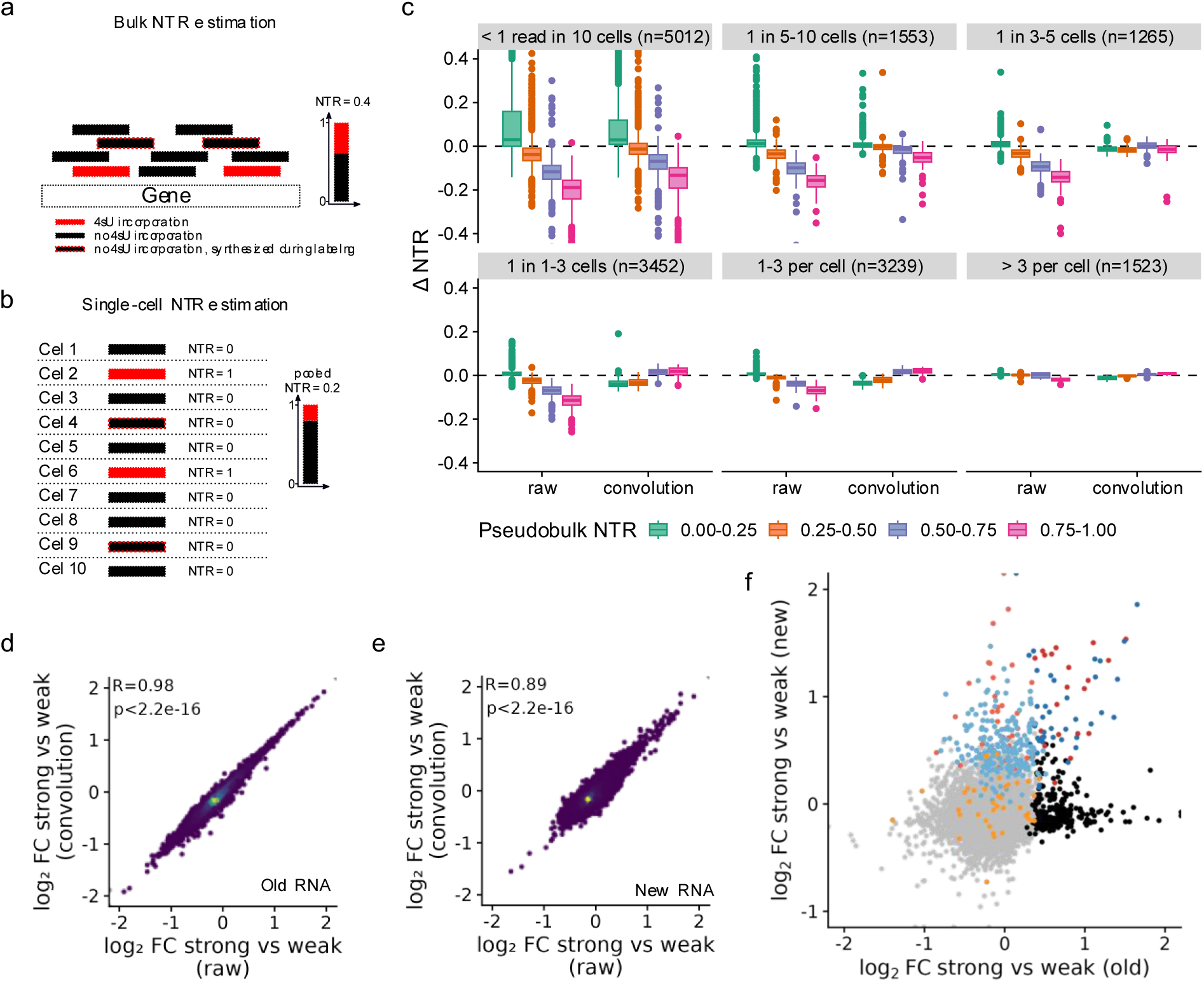
NTR convolution. **a** Illustration of bulk NTR estimation. Several reads (filled boxes) that originate either from RNA transcribed during labeling (red or red border) or not (black), and that either have 4sU incorporated (red) or not (filled black) are mapped to a gene. By its mixture model, GRAND-SLAM provides unbiased NTR estimates. **b** Illustration of single-cell NTR estimation. If the same reads shown in **a** originate from separate cells, the NTR estimate is biased. **c** Comparison of NTR values without (“raw”) and after the NTR convolution approach (“convolution”) in the DEX-treated data set. Genes were grouped according to expression level (panels) and NTR (colors) as indicated. ΔNTR represents the difference of gene-wise NTR of either raw single cell or convoluted estimates to pseudobulk-based NTRs (Boxplots: center line, median; box limits, upper and lower quartiles; whiskers, 1.5x interquartile range; points, outliers). **d** Scatterplot comparing log_2_ fold changes of pseudobulks of strong (top third DEX score) versus weak (bottom third DEX score) responder cells at the old RNA level before (“raw”) and after convolution. **e** Scatterplot comparing log_2_ fold changes of pseudobulks of strong (top third DEX score) versus weak (bottom third DEX score) responder cells at the new RNA level before (“raw”) and after convolution. **f** Scatterplot comparing log_2_ fold changes of pseudobulks of 2h cells versus 0h cells at the old RNA level to the new RNA level after NTR convolution. The same gene stratification as in Fig. 1g is used.

To mimic the beneficial situation in bulk RNA-seq, where many reads are available for most genes, we reasoned that information for more precise NTR estimates could be borrowed from cells that are close in the gene expression manifold. Thus, for each cell, we aggregated reads from its nearest neighbors and included these for the estimation of the NTR. To compare these ’convolution NTRs’ to the standard single-cell NTRs, we again pooled new and old RNA estimated by the convolution NTRs from all 0h cells and compared them to the pseudo-bulk NTRs. There were still large differences for genes that were expressed with less than 1 detected copy per 10 cells (n=5,012, 31.2%). However, for most genes, the single-cell convoluted NTRs were largely consistent with the pseudo-bulk NTRs (**Fig. 2c**).

We thus computed convoluted NTRs for all genes and repeated the analysis comparing old versus new RNA fold changes of strong versus weak responder genes. Of note, neither old (R=0.98, **Fig. 2d**) nor new RNA (R=0.89, **Fig. 2e**) fold changes were substantially affected by convolution on the pseudo-bulk level. As a consequence, the genes that appeared to be regulated on both the new and old RNA level remained largely unchanged (**Fig. 2f**).

We concluded that while the convolution approach provides more accurate estimates of the NTR at the single-cell level, it did not correct a potential spillover of new RNA in the old RNA.

### Single-cell trajectories connect pre- to post-treatment cells

As old RNA approximates a previous gene expression state of a cell, it can be used to connect cells to predecessor cells from a previous time point, and also across multiple time points to form single-cell trajectories ^15^. Each single cell trajectory is a sequence of cells from subsequent time points and reflects the progression of an individual cell over time. Such trajectories can be exploited for Heterogeneity-seq: The last time point provides the read-out of the outcome; the first time point represents the state of the cell before the stimulus.

A previously published method computed single cell trajectories by (***i***) approximating the previous transcriptome state for each cell by taking into account RNA degradation in between the time points, (***ii***) using canonical correlation analysis for each pair of subsequent time points to jointly embed the previous transcriptome state and the previous time point, and (***iii***) linking cells to the closest match either forward or backward in time ^15^.

Indeed, the UMAP of the joint embedding from the first two time points of the DEX time course indicated that the 2h cell population reflected the 0h cell population when using 2h previous RNA (degradation corrected unlabeled RNA) and 0h total RNA (**Fig. 3a**). However, when linking 2h cells to the closest match at 0h, only 627 out of 1,054 (59.5%) 0h cells were connected to 2h cells (**Fig. 3a, inlay i**). This is because the same 0h cell can be the closest match of multiple 2h cells. 271 of the 627 cells (43.22%) were connected to more than one 2h cell, and 163 of these 271 cells (60.15%) were connected to cells with multiple DEX score response groups (low, middle, high). Furthermore, when computing longer trajectories over multiple time points by this ’closest match’ approach and starting from the last time point, successively more and more cells were disconnected (**Fig. 3b-c**). In the full DEX treatment time course involving 6 time points, only 160 out of 1054 (15.2%) 0h cells were connected by single-cell trajectories. Taken together, the closest match approach cannot be used for Heterogeneity-seq, since rather few cells remain that do not cover the full spectrum of the first time point.

**Figure 3:**
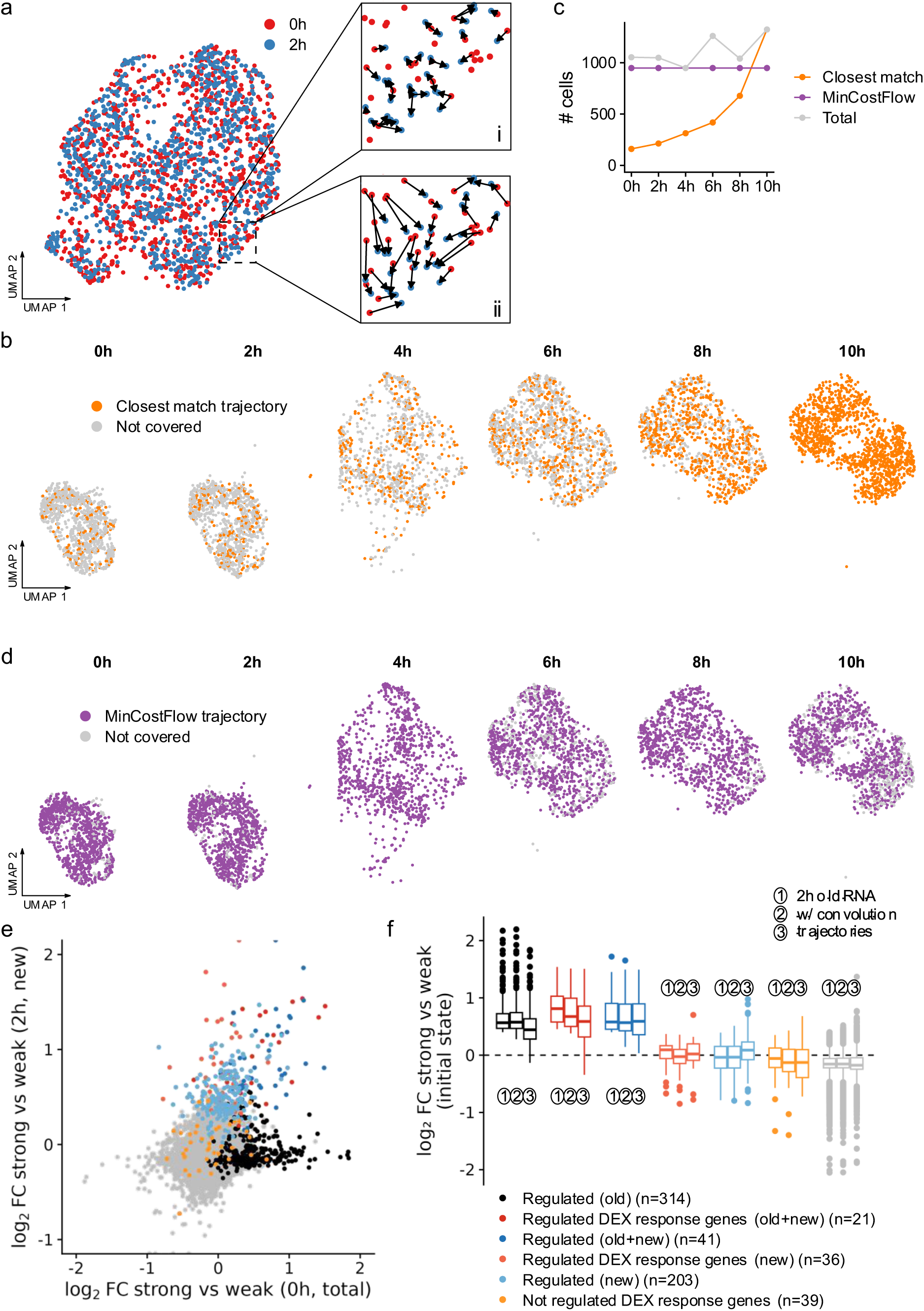
MinCostFlow trajectories in the DEX treatment sci-fate data. **a** UMAP representation of jointly embedded (CCA) 0h (total RNA) and 2h (previous RNA) cells. Inlay i shows trajectories (0h → 2h) constructed by matching each 2h cell to its closest 0h cell (“closest match”). Inlay ii shows trajectories constructed using MinCostFlow. **b** UMAP representation split by time point. Cells that are part of a trajectory constructed by the closest match algorithm are indicated. **c** Lineplot showing the number of cells (# cell) that are part of a trajectory constructed with either the closest match or MinCostFlow algorithms, as indicated, by time point. Also the total number of cells per time points are shown. **d** UMAP representation split by time point. Cells that are part of a trajectory constructed by the MinCostFlow algorithm are indicated. **e** Scatterplot comparing log_2_ fold changes of pseudobulks of 2h cells (new RNA) versus 0h cells (total RNA) that were connected via trajectories. The same gene stratification as in Fig. 1g and Fig. 2f is used. **f** Boxplot comparing the so-far proposed measures of the initial state of cells: 2h old RNA, x axis in Fig. 1g; 2h old RNA upon convolution, x axis in Fig. 2f; total RNA of 0h cells connected via trajectories, x axis in Fig. 3e per gene stratum (Boxplots: center line, median; box limits, upper and lower quartiles; whiskers, 1.5x interquartile range; points, outliers).

Thus, inspired by methods based on optimal transport theory ^26^, we developed an algorithm to compute optimal non-overlapping single-cell trajectories. A set of trajectories is non-overlapping if there is no cell that occurs in more than one trajectory. It is optimal if it has the minimal total distance among all sets of non-overlapping trajectories (see Methods). The problem of finding the optimal non-overlapping set of single-cell trajectories can be reduced to the minimum cost flow (MinCostFlow) problem and can therefore be solved efficiently using a linear program (**Fig. 3a, inlay ii**). The number of trajectories is by design equal to the minimal number of cells at a time point, and for the full DEX time course, trajectories fully cover the cell populations at each time point (**Fig. 3c-d**). In summary, similar to optimal transport, optimal non-overlapping single cell trajectories compute cell couplings among time points with the goal to cover the entire population from all time points. The main difference is that for optimal non-overlapping single-cell trajectories, the couplings are one-to-one mappings, whereas for optimal transport, they represent a joint probability distribution.

We next used this algorithm to implement Heterogeneity-seq. We computed short trajectories over the 0h and 2h time points, and, instead of comparing old and new RNA from the 2h time point (see **Fig. 1g and 2f**), we now compared new RNA from the 2h time point to total RNA from the connected 0h (mock) cells (**Fig. 3e**). Remarkably, the overall picture did not change: Overall, genes that were regulated at old RNA level between weak and strong responders were also regulated in the 0h cells connected via the trajectories to weak or strong responders, and genes that were not regulated stayed not regulated (**Fig. 3f**). Of note, also the genes that were regulated both at new and old RNA level were still regulated in connected 0h cells, suggesting that they are indeed both regulated and HetSeq factors, and that spillover of new to old RNA does not play a role. To further scrutinize trajectories, we recomputed trajectories after excluding all regulated genes from the distance metric. Still, genes regulated at both new and old RNA levels were also regulated at the 0h time point (**Extended Data Fig. 1**).

We concluded that the MinCostFlow algorithm can reconstruct faithful single-cell trajectories that reflect the likely temporal progression of single cells, and that there are indeed many genes that were both regulated and genes that are HetSeq factors for the DEX time course experiment.

### Double machine learning for inference of causal factors

HetSeq factors identified as differentially expressed before treatment in weak and strong response groups are associated with the response. However, whether they have direct effects (gene A causally affects the outcome), indirect effects (gene A has an effect on gene B, and gene B causally affects the outcome), or whether there is a confounding factor (gene B has an effect on both gene A and the outcome) is unclear.

Finding HetSeq factors can be phrased as a classification problem: Given the pre-treatment expression values of a candidate factor as a feature, the task is to predict the outcome (weak versus strong). If the outcome can be predicted based on a candidate gene, it is a HetSeq factor. We implemented this approach using a support vector machine (SVM), cross-validation, and the area under the receiver operating characteristic curve (AUC) as a measure of prediction performance. The advantage of this approach is that factors that are likely confounders can be integrated as additional features, and the lack of improvement in predictive performance can be used for excluding factors that only correlate with the confounder, but do not affect the outcome. Furthermore, we implemented double machine learning (dML) to systematically exclude confounding factors: In this approach, a model predicting the expression of a candidate factor from the expression of all other genes and a model predicting the outcome from all other genes are learned. In principle, this regresses out the influence of possible confounders from both the outcome and the candidate factors. If, after regressing out confounders, the outcome can be predicted from the candidate, it is considered a potential causal factor. This approach also allows us to estimate P values using influence functions and asymptotic theory ^27^.

We tested the extent to which correlations among genes from the 0h time point affect results, and how well causal inference can identify truly causal factors. Performing *in silico* simulations, we first binned genes into 10 groups according to their correlation with other genes. Group 1 consisted of genes that had the weakest correlations to all other genes, and genes in group 10 were strongly correlated with other genes (**Extended Data Fig. 2**). Of note, the average expression strength also increased from group 1 to 10 (**Extended Data Fig. 3**). We repeatedly selected 5 random genes from each group, defined these to be causal factors for the simulation and analyzed their ranks after performing Heterogeneity-seq using the SVM and dML. For the SVM, the 5 causal genes were ranked among the top 10% of all genes in 92.3% of all simulations for groups 2-8. For weak expression (group 1) and strong correlation with other genes (groups 9-10), the SVM performed less well (in top 10% in 64.7% of simulations, **Fig. 4a**). By contrast, dML based Heterogeneity-seq could in almost all cases identify the causal genes from other correlating genes: The 5 causal genes were the top 5 genes in 99.7% of the simulations for groups 2-8, and in 92.0% for groups 1, 9 and 10 (**Fig. 4b**).

**Figure 4:**
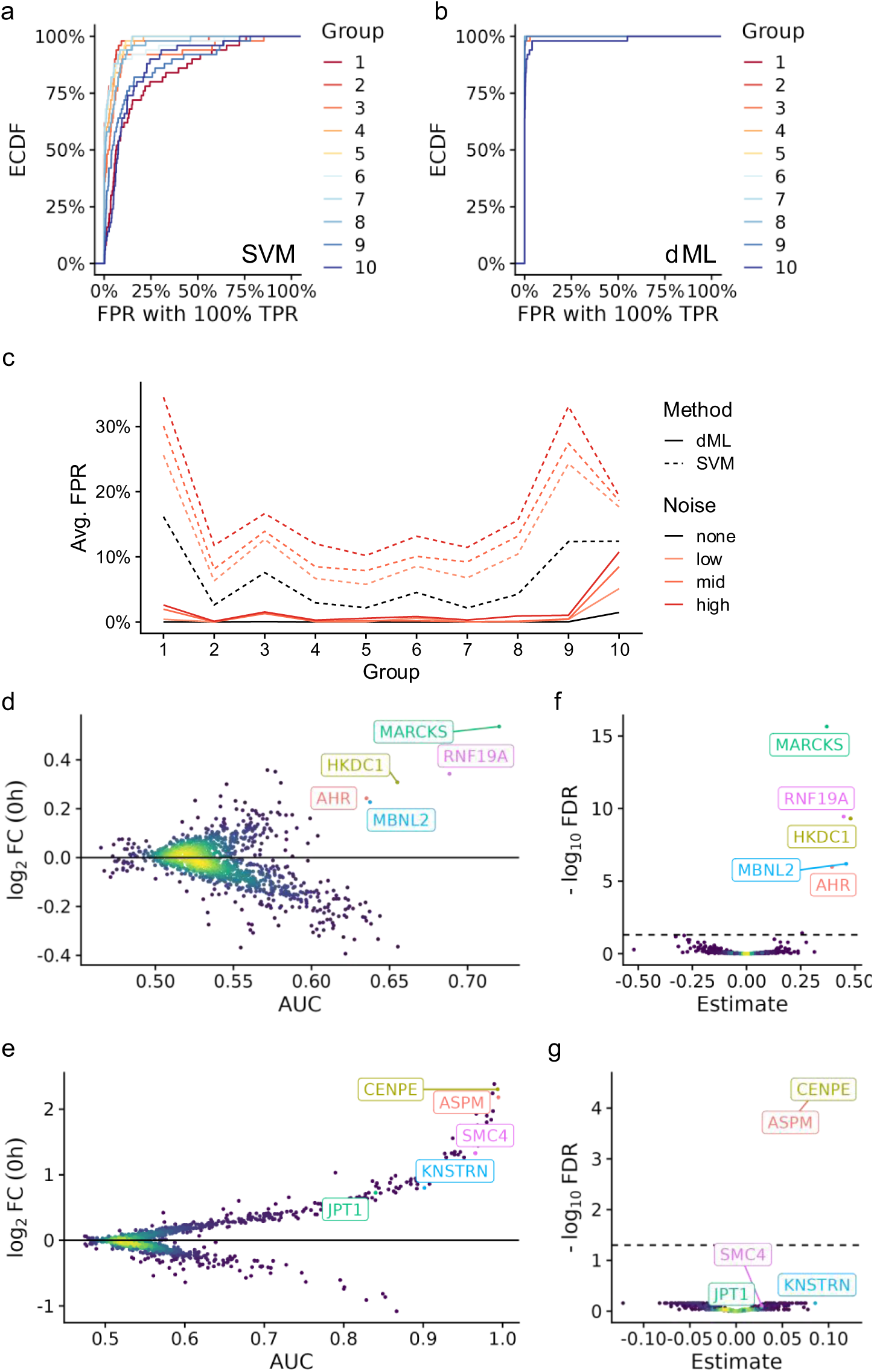
Causal inference validation by simulation. **a-b** Empirical cumulative distribution (ECDF) plot of the false positive rate (FPR) with 100% true positive rate (TPR) for the SVM-based (**a**) and double machine learning (dML, **b**) Heterogeneity-seq approach. The truth was repeatedly simulated for random subsets of genes selected from the indicated groups. For each simulation, the most stringent cutoff providing 100% TPR was chosen. **c** Lineplot showing the average FPR at 100% TPR (computed as in Fig. 4a**-b**) for both approaches (SVM and double machine learning) with different noise levels in the simulation. **d-e** Classification-HetSeq plot showing for each tested gene the area under the receiver operating curve (AUC) and the average log2 fold change at the 0h cells (low versus strong responders) for the SVM-based approach. The 5 genes (for **d**: group 3, for **e**: group10) defined to be causal factors in this simulation are indicated. **f-g** DoubleML-HetSeq plot showing for each tested gene the effect size (Estimate) and the multiple testing-adjusted (Benjamini-Hochberg) p value for the double machine learning approach. The 5 genes (for **f**: group 3, for **g**: group 10) defined to be causal factors in this simulation are indicated.

We also tested the impact of noise introduced by trajectories by randomly perturbing the distance matrix used to build trajectories. We prepared three noise levels, where 27.1% (low noise level), 34.4% (mid noise level) or 43.0% (high noise level) of all trajectories were rewired, and again analyzed the false positive rate when predicting all 5 causal genes (FPR_100_). The average FPR_100_ among repeated simulations increased from 3.6% (original trajectories) to 8.2% (low noise level), 10.1% (mid noise level) and 13.0% (high noise level) for the SVM approach in groups 2-8 and exceeded 30% for groups 1, 9 and 10 with high noise levels (**Fig. 4c**). Again, by contrast, for dML, the performance was near perfect also for high noise levels for groups 2-10 (average FPR_100_ below 0.7%), and only declined for group 10 (10% average FPR_100_).

To further investigate this drop in performance, we analyzed individual examples (**Fig. 4d-g**). While the gene combination *MBNL2, MARCKS, RNF19A, AHR,* and *HKDC1* from group 3 performed well (FDR_100_=0.4%) for the SVM (**Fig. 4d**), and perfect for dML (FDR_100_=0, **Fig. 4e**), the gene combination *KNSTRN, ASPM, SMC4, CENPE,* and *JPT1* from group 10 had an FDR_100_ of 6.0% for the SVM and 19.7% for dML (**Fig. 4f-g**). For this combination, dML only predicted *CENPE* and *ASPM* to be causal factors, presumably because of strong correlations among these 5 genes (all R>0.31).

Taken together, our simulations demonstrate that both the classification and double machine learning based Heterogeneity-seq approaches are effective in identifying causal factors, even when nearly half of all trajectories are incorrect. The SVM-based method consistently ranks causal factors highly, along with sets of other strongly correlating factors. The double machine learning approach is robust, accurately identifying and separating causal from correlating factors, but can miss causal factors when they correlate strongly. We concluded that both methods can be utilized to predict causal factors, avoiding potentially false positive hits (double machine learning) or a more lenient set of candidate genes, including directly causal genes (classification).

### Heterogeneity-seq recovers known factors impacting DEX responses

Having established an implementation of Heterogeneity-seq using optimal single-cell trajectories and classification or double machine learning predicting outcomes from the endpoints of trajectories using features from their starting points, we first investigated factors that impact the strength and quality of the response to stimulating the glucocorticoid receptor in A549 cells.

We first defined cell subsets with strong and weak responses in the 2h sample as above and applied Heterogeneity-seq. The basic approach of detecting differential expression of 0h cells connected to strong or weak responder 2h cells found 464 and 571 genes significantly contributing to a weak and strong DEX response, respectively (**Fig. 5a**, **Supplementary Table 1**). Upon further inspection with GO analysis, we found that the majority of genes negatively impacting the DEX response is connected to mitosis (**Supplementary Table 2**), which is in line with a study that showed a significantly lowered strength of the DEX response during mitosis and enforcing the idea of a novel function of steroid hormone receptors ^28,29^. The gene candidates related to a higher DEX response are associated with a wider variety of GO categories, e.g., wound healing, cell migration, and MHC complex assembly (**Supplementary Table 3**). Both results indicate that ‘cell cycle’ could be a confounding factor. We therefore used cell-cycle time (**Extended Data Fig. 4**, Methods) as an additional feature in our SVM and double machine learning based Heterogeneity-seq methods to identify genes independent of cell cycle-related effects. Among the best predictors in the classification model (**Fig. 5b**, AUC > 0.77, **Supplementary Table 4**), which were confirmed by causal inference (**Fig. 5c**, Benjamini-Hochberg corrected p < 0.05, **Supplementary Table 5**) were genes directly linked to the nuclear receptor metapathway or part of cytoskeletal signaling, which is of major importance for the transcriptional effect of the glucocorticoid receptor ^30^. Several of the top positive hits like *TRAM1*, *ZFP36L1* or *ACTN4* (**Fig. 5d**) are known to enhance glucocorticoid signaling ^31–33^. In addition, *SRGN* is known to be involved in resistance to a range of drugs in cancer ^34^. *G6PD*, which confers resistance to glucocorticoids was detected by classification but not the more restrictive double machine learning model ^35^. Conversely, double machine learning reported *XPO1* as additional negative modulators, which is known to play a major role in drug resistance ^36^. We concluded that Heterogeneity-seq can identify bona-fide causal factors that play a role in the extent of the early (2h) DEX response in single cells.

**Figure 5:**
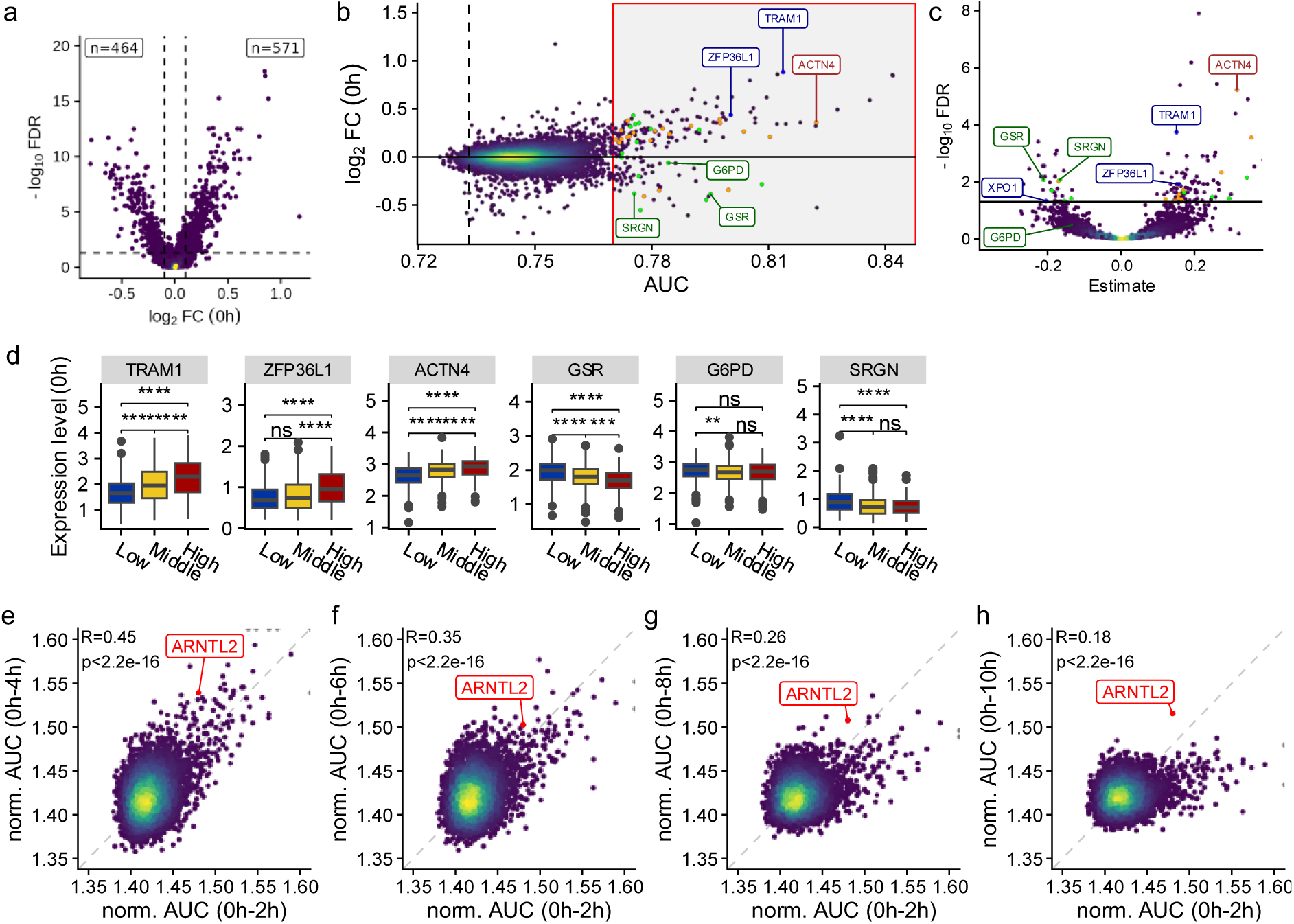
Heterogeneity sequencing for factors governing DEX responses in A549 cells. **a-c** 2h cells were stratified into low, middle, and high according to their DEX score, and MinCostFlow trajectories were computed connecting 0h cells to 2h cells. Data points are colored by point density. **a** Vulcano plot showing results of the differential expression approach to Heterogeneity-seq comparing the log_2_ fold change and Wilcoxon p values of pseudobulks of 0h cells connect the high versus low stratum. **b** Scatter plot comparing the log_2_ fold change to the predictive value (area under the receiver operating curve - AUC) for each gene. The AUC was computed for each gene by classifying cells as low or high, utilizing the 0h expression value and the cell cycle as features. The red box represents genes with AUC > 0.77. Specific gene sets were highlighted (cytoskeletal signaling, orange; nuclear receptor metapathway, green; manual curation, red). The dashed line represents the AUC value of cell cycle information only. **c** Vulcano plots of double machine learning model results, estimating the gene-wise effect on high or low DEX score outcome. **d** Boxplots of total RNA gene expression (0h) of highlighted genes, stratified by response groups in 2h, without taking cell cycle information into account (center line, median; box limits, upper and lower quartiles; whiskers, 1.5x interquartile range; points, outliers). Statistical significance is indicated (Wilcoxon test, ns, p > 0.05; *, p < 0.05; **, p < 0.01; ***, p < 0.001; ****, p < 0.0001). **e-h** Comparison of SVM-based Heterogeneity-seq across all time points (4h, 6h, 8h, 10h) versus 2h. For each run, response groups were defined on either 2h, 4h, 6h, 8h, or 10h cells (top third versus bottom third), trajectories were computed and classification was performed using 0h expression values including cell cycle as additional feature. Displayed are AUC values for all tested genes (n=7034 genes) normalized by the sum of all AUC values per time point and multiplied by 10,000. For all plots, Pearson correlation and p-values (two-sided approximate t-test) are indicated.

Next, we used Heterogeneity-seq to find factors where weak or strong expression before stimulation impacted the response strength at later time points (**Fig. 5e-h**). The impact on the response at 2h correlated moderately with the impact at 4h (R = 0.45), and a small set of genes (n=54) was detected at 2h but progressively lost predictive power, reflected in their AUC values decreasing over time. This observation could be due to additional noise introduced by the longer trajectories. However, we could exclude this because adding artificial noise to the 2h trajectories still maintained this set of genes (**Extended Data Fig. 5**). This indicates that these genes influence how fast the DEX response occurs in a single cell rather than how strong the response is. Interestingly, the top hit at 10h is *ARNTL2*, which is consistently among the top-ranked genes and known to be a key regulator of the circadian rhythm, which itself is inherently linked to the glucocorticoid system ^37,38^. This suggests that *ARNTL2* indeed impacts the strength of the response, in contrast to all other factors found. Taken together, these analyses demonstrate that by using single-cell trajectories, Heterogeneity-seq can also be applied over longer time frames to predict causal factors and also their mode of action.

### Heterogeneity-seq identifies novel pro- and antiviral host factors for cytomegalovirus infection

Next, we applied Heterogeneity-seq to the first two hours of murine cytomegalovirus (MCMV) infection. To this end, we first established a protocol for scSLAM-seq on the 10x Chromium platform (see methods). We sequenced overall 2,239 cells (Mock or MCMV infected) with a median UMI count of 35,832. In the UMAP and via clustering based on new RNA, cells segregated into three groups (**Extended Data Fig. 6a**) showing highly heterogeneous viral RNA load of up to 13.5% viral RNA (**Fig. 6a**). Accordingly, clusters were labeled mock, bystander, and infected cells, and infected cells were further subdivided into cells with low and high viral load (LVL/HVL) (**Fig. 6b**). The mock cluster (n=955, 42.7% of all cells), as expected for virus naïve cells, was characterized by minuscule levels of viral RNA (0.02%), presumably corresponding to ambient RNA (**Fig. 6c**). In bystander cells (n=646, 28.9% of all cells), 0.08% of all detected RNAs were viral, which was significantly more than in mock cells (p=7.0x10^-130^, two-sided Wilcoxon test). Moreover, bystander cells clearly induced *Ccl2* at levels like infected cells (**Figure 6d-e**). Ccl2 is a non-canonical interferon stimulated gene that is regulated by Irf3 downstream of pattern recognition receptors ^39^ (**Extended Data Fig. 6b**). Since we did not detect expression of any interferon gene, the induction of Ccl2 is likely because of cells sensing viral associated molecular patterns. Thus, bystander cells were indeed infected by MCMV, but the viral gene expression program has not been established. In LVL (n=384, 17.2% of all cells) cells 0.94% of all mRNAs were viral, whereas the viral RNA load was 3.8% in HVL cells (n=254, 11.3% of all cells). We concluded that the expression data enabled us to clearly differentiate mock, bystander, LVL and HVL cells, and that viral gene expression was highly heterogeneous among all infected cells.

**Figure 6:**
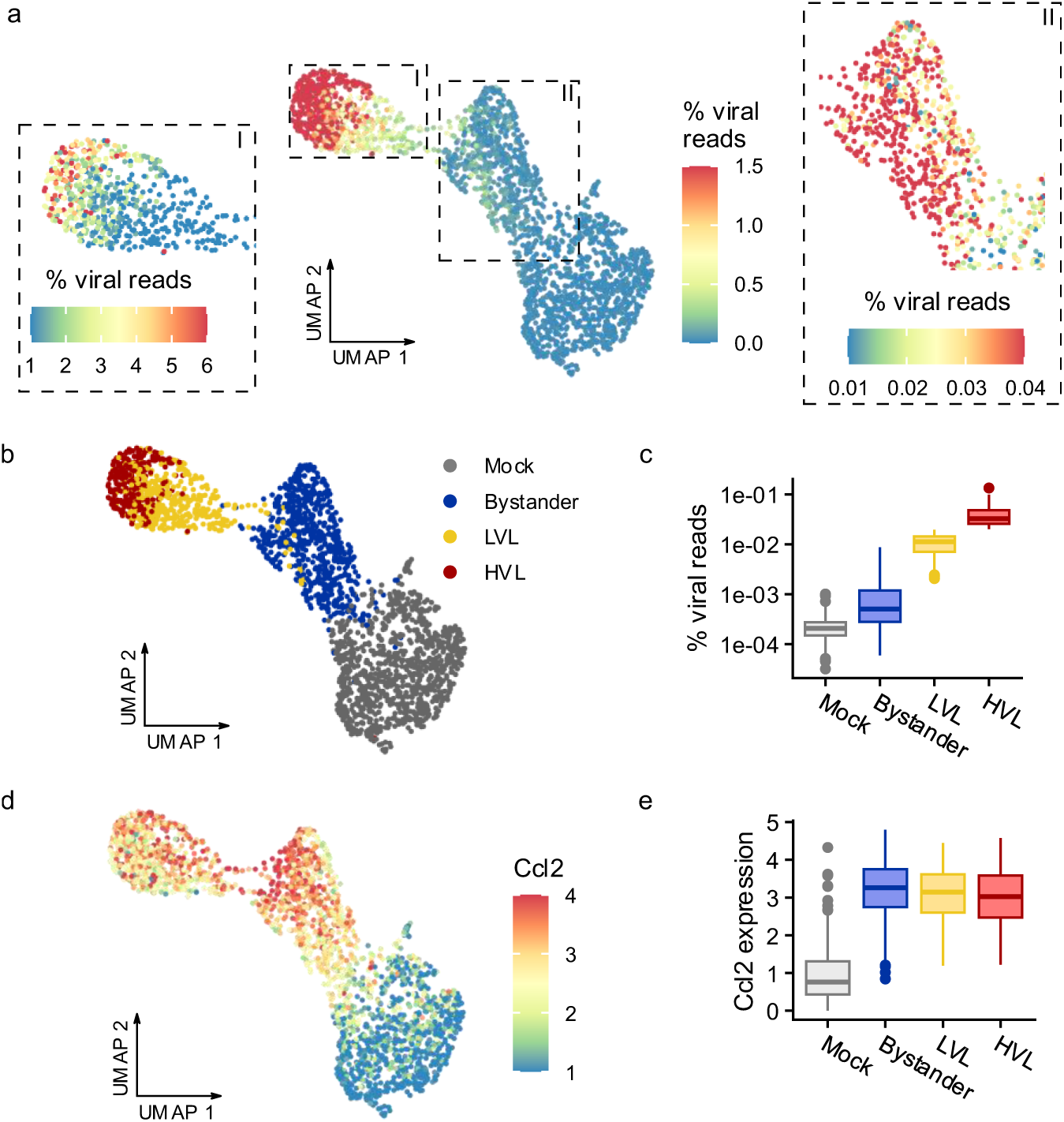
scSLAM-seq of MCMV infection in NIH-3T3 cells. **a** UMAP embedding of n=2,239 NIH-3T3 mock or murine cytomegalovirus infected cells. Cells are colored according to their viral RNA load (% of viral RNA). The two inlays I and II show zoomed-in cutouts, as indicated, using different color scales. **b** The same UMAP representation as in **a**, but with cells colored and labeled as mock, bystander and LVL/HLV according to the unsupervised clustering (Extended Data Figure 6) with the only exception that the upper cluster was split into two according to viral RNA load (<2%, low viral load LVL; >2%, high viral load, HLV). **<u>c</u>** Boxplots (center line, median; box limits, upper and lower quartiles; whiskers, 1.5x interquartile range; points, outliers) comparing the distributions of viral RNA load among the four clusters. **d** The same UMAP representation as in **a**, but with cells color by expression of Ccl2. **<u>e</u>** Boxplots (center line, median; box limits, upper and lower quartiles; whiskers, 1.5x interquartile range; points, outliers) comparing the distributions Ccl2 expression among the four clusters.

To investigate causal factors that govern these heterogeneous infection outcomes, we performed Heterogeneity-seq comparing HVL and bystander cells. To exclude impact of the cell cycle, we excluded all non-G1 phase cells from the analysis (**Extended Data Fig. 7**) and included the cell cycle time in our classification and causal inference system (**Figure 7**, **Supplementary Table 6-7**). Among the top factors predicted to facilitate the first two hours of infection (pro-viral host factors) found by classification (AUC > 0.715), we found genes connected to the microtubule or actin cytoskeleton like *Acta2* and *Tubb4b* (**Figure 6F**), which play a role in the transport of virus particles towards the nucleus ^40,41^. We also found tetraspanins *Tspan4* and *Cd63*, which are important parts of exosomes that are hijacked by herpesviruses for virion maturation, as well as the HCMV entry receptor Itgb1 ^42,43^. *Orc2* is a component of the origin recognition complex, playing a crucial role in DNA replication ^44^. Conversely, *Orc2* depletion has been shown to enhance HCMV DNA replication ^45^. The anti-viral hits mostly comprise genes related to ribosomes and the process of ribosome biogenesis, which itself seems to facilitate dsDNA sensing and confer antiviral properties ^46^. Also, the intermediate filament protein *Vimentin* was predicted to be anti-viral by classification, which has been reported to have both pro- and anti-viral properties ^47^. Our double machine learning model confirmed *Orc2*, the actin-related genes *Acta2* and *Myl6* as well as the microtubule related genes *Kif1c* and *Tubb4b*, whereas the tetraspanins *Tspan4* and *Cd63* failed to reach statistical significance. We concluded that the SVM-based approach was able to clearly identify broad classes of genes that either facilitate (actin, microtubule, tetraspanin) or block (ribosome related) viral gene expression in this very early phase of infection. By contrast, the double ML approach identifies much fewer genes.

**Figure 7:**
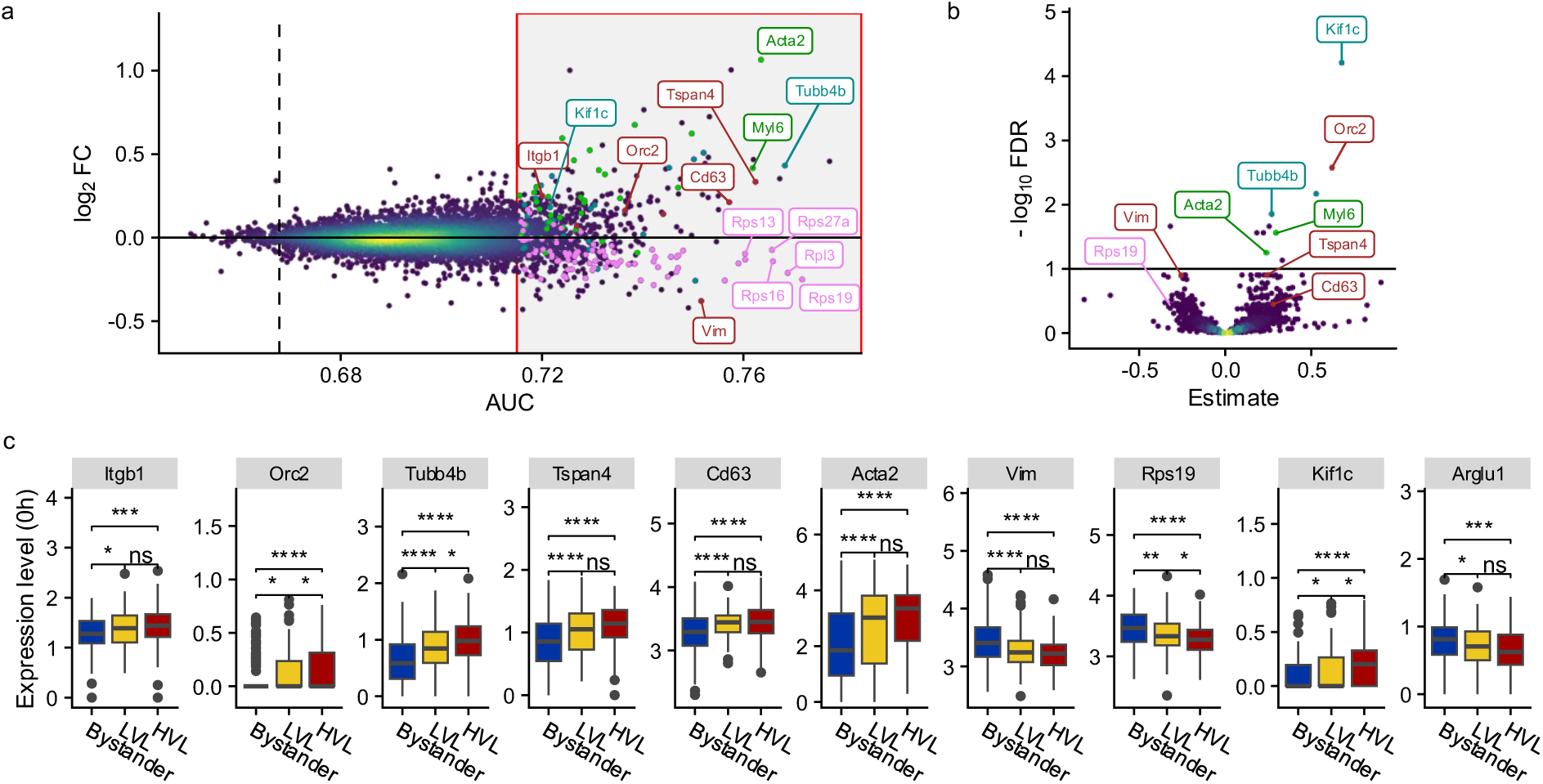
Heterogeneity-seq for factors governing MCMV infection outcomes in NIH-3T3 cells. **a** Scatter-plot comparing the log_2_ fold change to the predictive value (area under the receiver operating curve - AUC) for each gene. The AUC was computed for each gene by classifying cells as Bystander or HVL utilizing the 0h expression value and the cell cycle as features. The red box represents genes with AUC > 0.715. Specific gene sets were highlighted (microtubules, cyan; actin, green; ribosomes, violet; manual curation, brown). The dashed line represents the AUC value of cell cycle information only. **b** Vulcano plot of double machine learning model results, estimating the gene-wise effect on Bystander or HVL. **c** Boxplots of total RNA gene expression (0h) of highlighted genes, stratified by response groups in 2hpi, without taking cell cycle information into account (center line, median; box limits, upper and lower quartiles; whiskers, 1.5x interquartile range; points, outliers). Statistical significance is indicated (Wilcoxon test, ns, p > 0.05; *, p < 0.05; **, p < 0.01; ***, p < 0.001; ****, p < 0.0001).

## Discussion

Heterogeneity-seq provides a framework for identifying causal molecular factors that govern cell-to-cell variability in the acute response to molecular challenges such as drug treatment or viral infection. By exploiting naturally occurring transcriptional heterogeneity and linking pre-stimulus states to post-stimulus outcomes via metabolic RNA labeling, Heterogeneity-seq goes beyond purely correlative single-cell analyses towards identifying truly causal factors. In contrast to active perturbation screens, no genetic manipulation is required, making it applicable to virtually any biological system.

A misleading strategy to identify causal factors would be to focus on genes that are differentially expressed after stimulation. However, expression differences might arise because of differential expression before the stimulus (in which case it might be a causal factor), or because they are themselves regulated as part of the response (in which case it likely is no causal factor), or both. These scenarios are indistinguishable when only analyzing post-stimulus cells, and make the directionality of causality ambiguous. Taking all regulated genes as causal would inflate false positives, while removing them altogether would mask true factors that are also transcriptionally responsive. Heterogeneity-seq addresses this ambiguity by explicitly linking pre-stimulus and post-stimulus states of individual cells through metabolic labeling and trajectory inference, allowing the identification of genes whose prior expression predicts subsequent molecular outcomes.

Despite advances in quantifying labeled and unlabeled RNA and removing bias ^17,48^, the separation between these two RNA pools is not perfect. The limited incorporation rate of 4sU and short read length can lead to contamination between labeled and unlabeled fractions. Reads originating from newly synthesized transcripts may lack T-to-C conversions and thus be misclassified as pre-existing RNA, while sequencing or amplification errors can create spurious conversions in unlabeled RNA. Both effects blur the distinction between old and new RNA, particularly for single-cell experiments due to overall small read numbers. The convolution approach implemented here mitigates some of these issues by borrowing information from neighboring cells, yet it cannot fully eliminate them.

A conceptual limitation of Heterogeneity-seq lies in the reconstruction of single-cell trajectories. The trajectories are, by design, inferred between different cells sampled at successive time points. However, the same cell is not followed physically via sequencing. Thus, any trajectory is necessarily an approximation. However, if we view the transcriptomes of cells as points on a high-dimensional manifold representing the continuum of cell states and consider how cells would move upon the external molecular challenge, the trajectories effectively connect pre- and post-stimulus regions. Using unlabeled RNA instead of total RNA to infer connections is essential in settings with rapid transcriptional transitions, such as acute stimulation or viral infection. The successful recovery of biologically meaningful factors in both drug treatment and infection contexts strongly suggests that these inferred trajectories provide a faithful coarse-grained representation of underlying cellular progressions.

The SVM-based classification approach is sensitive and can readily detect predictive relationships between pre-stimulus expression and outcome, but it cannot distinguish whether a gene is truly causal or merely correlated through a confounder. Although in principle any gene can be tested as a confounder by including it as a feature for the SVM, it does not provide a systematic way to exclude confounding factors. By contrast, the double machine learning (dML) framework explicitly accounts for potential confounders by regressing out shared influences from both predictors and outcome. This improves specificity but comes at a cost: When two or more genes are strongly correlated with each other before the stimulus, and one of them is indeed a causal factor, then all residuals might be noisy to an extent such that dML may be unable to assign statistically significant causality to any single member of the group. Then, even though from the data it is clear that at least one is very likely to be a causal factor, dML cannot detect which one it is and report none of them as significant. Thus, in practice, the two approaches offer complementary perspectives, SVM identifying broader sets of candidate regulators, and dML isolating a smaller but more confident subset of direct causal factors.

Experimental validation of candidate factors remains difficult. The Heterogeneity-seq concept avoids active perturbations to capture physiological variability, yet mechanistic validation ultimately requires knockouts or overexpression. Such active perturbations are done a relatively long time before the molecular challenge (typically days) and can therefore easily alter the very network topology that gave rise to the inferred causal relationships. Thus, validation must be guided by orthogonal evidence such as temporal consistency, concordance across datasets, or perturbation results interpreted within the inferred causal framework, rather than relying on single-gene knockouts alone.

In summary, Heterogeneity-seq leverages natural expression variability, metabolic labeling, and advanced statistical inference to extract causal insights from single-cell data without active perturbation. While the approach faces challenges related to labeling fidelity and the intrinsic approximations of trajectory inference, its ability to recover known and novel regulators demonstrates that these limitations are not prohibitive. The combination of classification and dML-based inference provides complementary strengths: one favoring sensitivity, the other specificity. Together, they offer a powerful route to dissect how heterogeneous cell states shape molecular outcomes in dynamically changing biological systems.

## Online Methods

### Cell culture and infection

NIH-3T3 (ATCC CRL-1658) Swiss mouse embryonic fibroblasts were grown in DMEM supplemented with 100 IU/mL penicillin, 100µg/mL streptomycin, and 10% NCS (new-born calf serum). M2-10B4 (ATCC CRL-1972) fibroblasts were grown in RPMI-1640 supplemented with 100 IU/mL penicillin, 100µg/mL streptomycin, and 10% FCS (Fetal calf serum). BAC-derived MCMV Smith strain was utilized for scSLAM-seq experiments ^49^. Infected M2-10B4 cells and supernatants were harvested after >90% infection for virus purification, and titration of virus stocks was conducted by standard plaque assays on NIH-3T3 cells ^50^. Infection was conducted at an MOI of 10. All cells were cultured at 37 °C with 5% CO_2_.

### scSLAMseq methodology and sequencing

NIH-3T3 cells were infected with MCMV for 2h and at the same time labeled with 400 µM 4sU as described ^9^. Afterwards, cells were trypsinized using mild trypsin (diluted with PBS in a ratio of 1:1). Cells were pelleted at 350 g at 4 °C for 5 minutes. The cell pellet was gently resuspended in 1 ml of cold 1XPBS. Subsequently. 4 ml of ice-cold methanol (−20°C) was added to the cells dropwise using a vortex set at low speed. Fixed cells were incubated on ice for 15-20 minutes and were then aliquoted and stored at -80 °C freezer.

On the day of the experiment, fixed cells were taken out from the -80 °C freezer and were thawed on ice for 15 minutes. Afterward, cells were centrifuged down at 350 g for 10 min at 4 °C. Methanol was removed, and the cells were washed once with 3X SSC containing 0.04% BSA. Cells were rehydrated in 3X SSC containing 0.04% BSA at 4 °C for 30 min. Subsequently, cells were centrifuged at 350g for 10 min at 4 °C and resuspended in 100 ml of 3X SSC containing 0.04% BSA. 100 ml of 20 mM iodoacetamide (IAA) in DMSO was added to the cells (10 mM final concentration). Cells were incubated at 37 °C for 30 minutes in the dark. Finally, 20 ml 1M DTT was added to the reaction mix to quench the IAA and was kept on ice for 5 min. 1 ml of 3X SSC containing 0.04% BSA was added to the cells and centrifuged for 10 min at 500 g at 4 °C. Cells were washed once more with 3X SSC containing 0.04%BSA. Finally, cells were resuspended in 0.3X SSC containing 0.04% BSA. Cells were counted. Infected and non-infected cells were mixed in equal numbers and were processed for droplet generation.

3,000 cells were loaded in the Chromium Controller for partitioning single cells into nanoliter-scale Gel Bead-In-Emulsions (GEMs). Single Cell reagent kit v3 was used for reverse transcription, cDNA amplification, and library construction of the gene expression libraries (10x Genomics) following the detailed protocol provided by 10x Genomics. SimpliAmp ThermalCycler was used for amplification and incubation steps (Thermo Fisher). Libraries were quantified by Qubit TM 2.0 Fluorometer (ThermoFisher), and quality was checked using 2100 Bioanalyzer with High Sensitivity DNA kit (Agilent). Sequencing was performed in paired-end mode with S2 and NextSeq500 High-output flow cells (2x50 cycles) using NovaSeq 6000 and Nextseq500 sequencer (Illumina).

### SLAM-seq data processing

For the sci-fate (DEX treatment) dataset, we downloaded all 190 fastq files from SRA (Accession id SRX5848248). To facilitate further processing, we generated a unique 4-letter barcode for each of the 190 fastq files. This enabled us to concatenate all 190 fastq files (appending the 4-letter barcode + read 2 to the read identifier) and proceed with our standard processing pipeline for nucleotide conversion RNA-seq data ^9,17,18,48^. The concatenated fastq file was processed using cutadapt (version 3.5) ^51^ to remove poly-A tails (parameters -a AAAAAAAA --overlap=6) and then mapped to the human genome (Ensembl v90) using STAR (version 2.7.10b) ^52^ with parameters --outFilterMismatchNmax 20 --outFilterScoreMinOverLread 0.4 --outFilterMatchNminOverLread 0.4 --alignEndsType Extend5pOfReads12 --outSAMattributes nM MD NH. The bam file created by STAR was then converted to the CIT format using the Bam2CIT tool from the gedi toolkit (version 1.0.6a) with parameters -umi. Finally, we used our GRAND-SLAM software (version 3.0.4) ^17^ to compute read count and NTR matrices.

The MCMV data set was mapped using the cellranger (version 5.0.1) count pipeline with –expect-cells=2000 and otherwise default parameters. We used a combined reference genome of the human (Ensembl v90) and MCMV C5X genome ^49^. The bam file created by cellranger was then converted to the CIT format using the Bam2CIT tool from the gedi toolkit (version 1.0.6a) with parameters -umi. Finally, we used our GRAND-SLAM software (version 3.0.4) ^17^ to compute read count and NTR matrices.

### GRAND-SLAM 3.0

Since the original GRAND-SLAM version ^17^, and the release of GRAND-SLAM version 2.0 ^9^, we developed its 3^rd^ version. The following paragraphs describe all improvements of GRAND-SLAM 3.0: Clipping parameters for excluding extreme mismatch frequencies at the 5’ or 3’ end of reads can now be detected automatically (or set manually as previously). The automatic detection is done as follows: First, for each position in all reads that are later used for counting, relative frequencies of T-to-C mismatches or A-to-G mismatches (if a read is sense or antisense, respectively) are determined. The relative mismatch frequency for position *p* is the number of T-to-C mismatches or A-to-G mismatches observed at position *p* in a read divided by the total number of reads that map to a genomic C or G with their position *p*. This is done for all *k* samples of an experiment. Each position is thus represented by a *k*-dimensional vector of relative frequencies, and finding positions to exclude becomes an outlier detection problem. Thus, the *k*-long mean vector and *k* × *k* covariance matrix of all “inner” (excluding the first and last 20%) positions is computed. Based on that, the Mahalanobis distances for all *k*-dimensional vectors corresponding to the positions along the reads are determined, and a *χ*^2^ test is used to compute p values for each position testing whether a position is not an outlier. The maximal position among the first 25 positions with a Bonferroni-corrected p value < 1% is used for 5’ end clipping and, likewise, the minimal position among the last 10 positions for 3’ end clipping. Diagnostic plots are produced and were checked for both data sets (scifate: 15 positions clipped at 5’ end, 3 positions at 3’ end, MCMV: 12 positions clipped at 5’ end, 2 positions at 3’ end).

GRAND-SLAM 3.0 provides two procedures to estimate global parameters: GRAND-SLAM is based on the mixture model 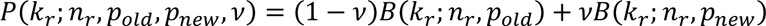 where 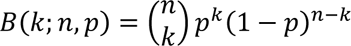 is the binomial probability, *k_r_* is the number of T-to-C mismatches observed for *n_r_* positions mapping to a genomic T in read *r*. The global parameters are the probability for a mismatch in a read from an unlabeled RNA *p_old_*, the probability for a mismatch in a read from a labeled RNA *p_new_* and the global new-to-total RNA ratio (NTR) *ν*. For the whole data set, *ν* is a nuisance parameter. Once the global parameters are found, the same model is applied to each gene: *p_old_* and *p_new_* are treated as fixed ^53^, and *ν* is estimated taking into account all reads mapped to the gene. The original implementation estimated *p_old_* from no4sU control samples, an *p_new_* with an EM algorithm from all reads with typically at least 3 T-to-C mismatches ^17^. The second procedure numerically maximizes the log likelihood function 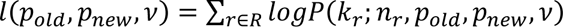 with *R* is the set of all reads compatible with a gene using the Nelder-Mead algorithm. For both the scifate and MCMV data sets analyzed here, the new procedure was used.

With GRAND-SLAM 2.0 ^9^, we introduced the idea that parts of a cDNA are sequenced more than once: For paired-end data, if the cDNA fragment that is sequenced is shorter than the total length of the read pair, the ends of both reads correspond to the same positions in the fragment (overlap part). For these positions, sequencing errors (but not reverse transcription errors or PCR errors) can virtually be excluded if they are not observed consistently in both reads. The same idea can also be used for data with UMIs. In GRAND-SLAM 3.0, each sequencing unit (e.g., a read, a read pair, or all reads from the same UMI) is separated into so-called subreads. A subread is a part of a sequencing unit where we expect uniform mismatch probabilities. E.g., for paired-end data, there are three subreads: (i) all positions only mapped by the first read, (ii) all positions only mapped by the second read, and (iii) all positions in the overlap part. For UMIs as produced by the 10x Chromium platform and also combinatorial indexing, we also defined three subreads as (i) all positions of a UMI only mapped by a single read, (ii) all positions mapped by exactly two reads and (iii) all positions mapped by three or more reads. Global parameters are estimated separately for each subread. Using the NTR *ν_g_* for a gene *g* requires one to compute the likelihood of all sequencing units for both components of the mixture model. If there are multiple subreads, this can be done by multiplying the binomial probabilities of all subreads *B*(*k*_1_; *n*_1_, *p*) ⋅ *B*(*k*_2_; *n*_2_, *p*) ⋅ … where *k_i_* is the number of T-to-C mismatches observed for subread *i* of the sequencing unit, and *n_i_* the number of positions mapping to a genomic T in subread *i*.

GRAND-SLAM 3.0 is a reimplementation with a completely new codebase which was necessary to make it applicable in a setting with thousands of cells. It is highly extensible, e.g. it can also be used for 6-thioguanosine (6sG) labeling and combined 4sU and 6sG labeling. It can work with any sequencing data (stranded/unstranded, single-end/paired-end, UMIs) and provides mechanisms to implement different estimation procedures and even different models. It can output results as standard table or as sparse matrices, both of which can be read by our grandR package ^18^.

### Quality control, filtering and general data analysis

Data was imported into R (version 4.3) from the GRAND-SLAM output using the grandR package (version 0.2.5). For the Scifate data set (7404 cells, 43750 genes), we filtered for the same cells as in the original publication (list of cells provided in the online material in ref. ^15^ and for genes that are expressed in at least 100 cells. After filtering, 6680 cells and 16046 genes remained. For the MCMV data set (2554 cells, 11113 genes), we filtered for cells with expression in at least 100 cells, less than 10% of mitochondrial genes in regard to total gene expression and a number of genes in 2000 < x < 8000. After filtering, the MCMV data set contained 2239 cells and 11020 genes.

Normalization (default parameters), data scaling (default parameters), dimensionality reduction (PCA and UMAP with 10 dimensions), Clustering (only performed on MCMV data set, resolution 0.1), general data analysis and visualization was performed using the Seurat (version 4.3.0) and ggplot (version 3.5.1) packages.

The viral load was calculated as the percentage of viral reads among all reads per cell. The DEX score was calculated from new RNA using the AddModuleScore function of the Seurat package, using a predefined set of dexamethasone response genes ^15^.

### Construction of optimal cellular trajectories

A trajectory is defined as a sequence of cells from consecutive time points of a single-cell time course. A set of trajectories is non-overlapping if each cell occurs at most once in a trajectory. A set of trajectories is optimal if (i) it is non-overlapping, (ii) each trajectory reflects a likely progression of a single cell over time, and (iii) the set is maximal (i.e., there are as many trajectories as possible in this set). Because of the first two conditions, each trajectory approximates a putative experiment, where a single cell is measured multiple times. The third condition ensures that the whole cell population under study is analyzed. To find likely progressions (ii), we leverage the information from metabolic RNA labeling, namely that unlabeled RNA reflects the cellular state from an earlier time point, whereas total RNA reflects the state of the cell at the current time point. A progression is considered likely if the previous state of a cell is similar to the current state of its predecessor in the trajectory.

More formally, to find an optimal set of trajectories, we require a distance function *d* for cells of subsequent time points, and we seek to find a set of trajectories which minimizes the total distance over all time points and trajectories among all non-overlapping, maximal sets of trajectories.

To find the optimal set of trajectories, we define the following flow network:

1. There is a source vertex *a* and a sink vertex *b*
2. _2._ For each cell *x* there are two vertices *p_x_* and *t_x_*
3. There are (directed) edges (*p_x_*, *t_x_*) for each cell *x*
4. For each pair of cells *x* and *x*′, where *x*′ is from the subsequent timepoint of *x*, there is an edge (*t_x_*, *p*_*x*′_).
5. In a similar manner, for each cell *x* from the last time point, there is an edge to the sink vertex (*x, b*), and for each cell *x*′ from the first time point, there is an edge from the source vertex (*a, x*^′^).
6. Each edge (*ν*, *u*) has a capacity *c*(*ν*, *u*) = 1.
7. Each edge (*ν*, *u*) defined under 3. or 5. has a cost of *a*(*ν*, *u*) = 0.
8. Each edge (*t_x_*, *p*_*x*′_) defined under 4. has a cost of *a*(*t_x_*, *p*_*x*′_) = *d*(*x, x*^′^) where *d* is the distance function

The network consists of layers of vertices: The first layer is the source vertex. The second and third layers consist of the *p*. and *t*. vertices of the first time point. The fourth and fifth layer likewise consist of the *p*. and *t*. vertices of the second time point etc. The last layer is the sink vertex. Edges only exist between two subsequent layers. For a *p*. layer to a *t*. layer, there are as many edges as there are cells in the time point. For all other pairs of subsequent layers, all vertices of the first are connected to all vertices of the second. A trajectory corresponds to a path from source to sink in the network.

For the following, we adopt the usual notation that a network flow is a function that assigns a flow (a real number) to each edge, such that the incoming flow is equal to the outgoing flow (flow conservation) for all vertices except for the source and the sink, and such that the flow for an edge is at most its capacity. The value of the flow is the sum of the flows of the edges outgoing from the sink.

**Lemma 1:** The maximum flow corresponds to a non-overlapping set of trajectories.

**Proof:** By the integrality theorem for maximal flows, the flow on each edge in the maximum flow is an integer (since all capacities are integer). Since the capacities are all 1, the maximum flow on each edge is therefore either 0 or 1. Now consider the network induced by all edges with flow 1. In this network each vertex except for source and sink has either zero neighbors, or exactly one predecessor and one successor. Because of flow conservation, any vertex connected to an edge must be connected to both source and sink via a path. Thus, this network consists of paths from source to sink, which correspond to non-overlapping trajectories.

**Lemma 2:** The value of the maximum flow of the network is equal to the number of cells in the bottleneck time point (i.e. the time point with the fewest cells) and therefore to a maximal and non-overlapping set of trajectories.

**Proof:** Let *n* be the number of cells in the bottleneck time point. Take a random subset of size *n* from each time point and arrange them randomly in a non-overlapping set of trajectories. According to the argument above, this non-overlapping set of trajectories is a network flow. Thus, there exists a network flow with value *n*. Since there exists a cut between the *p*. layer to the *t*. layer of the bottleneck time point with capacity *n*, according to the Max-Flow-Min-Cut-Theorem there cannot be a flow with value ≥ *n*.

**Theorem:** The minimum-cost flow in the network among all maximum flows corresponds to the optimal set of trajectories.

**Proof:** By Lemma 1 and 2, there is a one-to-one correspondence between maximum flow in the network and maximal, non-overlapping sets of trajectories. Therefore, conditions (i) and (iii) of an optimal set of trajectories are fulfilled for maximum flows. By construction, there is also a one-to-one correspondence between the cost of a maximum flow and the total distance of the corresponding trajectories. Let *T* be the set of trajectories corresponding to the maximum flow with minimal costs. If there was a set of trajectories *S* with lower total distance than *T*, there was a maximum flow corresponding to *S* with lower costs than *T*.

Thus, to find an optimal set of trajectories, we construct the flow network and solve the minimum-cost flow problem (which can be done using a linear program). To speed up the algorithm, we introduced a pruning step keeping only the top *k* incoming (according to their cost) edges for the *p*. vertices and top *k* outgoing edges for the *t*. vertices. According to our evaluations pruning with *k* = 10 did not change the trajectories.

To construct a distance function for cells from timepoints *n* and *n* + 1, we use the canonical correlation analysis integration implemented in Seurat: We use the standard workflow of normalization, variable feature identification, selection of integration features, finding of integration anchors and integrate on the count matrix of total RNA from time point *n* and on the count matrix of previous RNA (see below) from time point *n* + 1. We then compute the first 15 principal components on the integrated data and define the distance function as the squared Euclidean distance.

### Convolution-based NTR calculation and old/new read count recalculation

For the estimation of new-to-total ratios (NTRs), we used GRAND-SLAM ^17^, which, by default, collects all reads per cell for a specific gene to determine both the number of thymines in this genomic region as well as all T-to-C mismatches as a basis for the estimation. Our convolution-based approach adapts the GRAND-SLAM method by supplying a set of cells c_1_ … c_k_ for every cell c (i.e. the k-nearest neighbors of c), expanding the collection of reads to include all reads of cells c_1_ … c_k_ for every cell c as a new basis for the subsequent NTR estimation.

The convoluted data was read in as a grandR object and its table of NTRs was used to recalculate gene-specific new read counts by matrix multiplication of the NTR matrix with the total read count matrix of the non-convoluted data set. The old read counts were then determined by subtracting recalculated new read counts from the total read count matrix. These recalculated old and new RNA counts are then used for all subsequent analyses and generally referred to as old and new RNA counts.

### Calculation of previous RNA counts

To calculate previous RNA counts, reflecting the state of a cell at the beginning of the labeling perious, we first calculate half-lives using the ComputeSteadyStateHalfLives function of the grandR package on the convoluted grandR object and set minimum half-lives to 0.25h and maximum half-lives to 24h. We then calculate the previous RNA counts as follows:

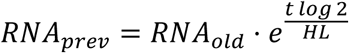

with *RNA_old_* representing old RNA counts, *t* representing the labeling time (2 hours for all data sets used in this study) and *HL* representing convolution-based half-lives.

### Cell cycle scoring

For the calculating of a cell cycle score, we use the convoluted data (see above). We import the data into R using the grandR package and transformed the convoluted grandR object into a Seurat object using as.Seurat.grandR(). Following the standard Seurat workflow, we performed data normalization using NormalizeData(), identified variable features with FindVariableFeatures(), and scaled the data using ScaleData(), all using default parameters. Principal Component Analysis (PCA) was conducted with RunPCA(). We constructed a K-nearest neighbor graph using FindNeighbors(), considering the first 10 principal components. Cell clusters were identified using the Louvain algorithm with FindClusters() (resolution = 0.2). Finally, we visualized the data using Uniform Manifold Approximation and Projection (UMAP) for dimensional reduction with RunUMAP(), again utilizing the first 10 principal components.

Following the UMAP visualization, we performed cell cycle scoring with Seurat’s CellCycleScoring() function, using the 2019 updated list of cell cycle markers from ref. ^54^. This function calculates scores for each cell based on the expression of genes known to be active during specific cell cycle phases (S-phase and G2/M-phase). We then conducted PCA (RunPCA()) using only these cell cycle genes, and used the first 3 principal components for clustering (FindClusters(), resolution = 0.2) and a new UMAP (RunUMAP()) visualization. This cell cycle-focused UMAP revealed a circular pattern, resembling their position in the cell cycle. From this, we can either take the first and second UMAP coordinates as cell cycle scores or create a singular score based on a cell’s position in the circle. To create this score for each cell, we implemented a method based on the angular position of cells in the UMAP space. We extracted the UMAP coordinates for all cells and calculated their geometric center. For each cell, we calculated its angle relative to this center using the atan2 function and the result was converted to degrees (0-360°). We then identified one cell as our starting point, which was positioned at the edge of G0-G1 phase and likely representing an early G1 state. All cell angles were adjusted relative to this starting point and normalized to a 0-1 scale by dividing by 360, creating our cell cycle score. We used the first and second UMAP coordinates as cell cycle scores in the Heterogeneity-seq analyses.

### In-silico simulation

To test the classification based and double machine learning Heterogeneity-seq approaches, we implemented an in-silico simulation algorithm: The simulation involves four major steps:

1. The calculation of a set of ground truth trajectories between 0h and 10h cells,
2. the definition of a response score for each cell in the 10h sample based on expression a defined set of genes in their predecessor cells (0h), and
3. the recalculation of trajectories with increasing noise levels.

To guarantee comparable results between simulations, we first restricted the available cells in the simulation to a fixed set of 949 cells per time point (based on the 4h time point which had the lowest number of cells). We then calculated a set of ground truth trajectories for the full time course (see Methods, Construction of optimal cellular trajectories).

These trajectories established the ground truth connection for each 0h cell to an endpoint cell from the 10h timepoint. The 10h cells are used to define a response score based on the expression of a specific set of genes in their direct predecessors. For that, we computed the module score of the given gene set in each 0h cell, and then transferred these scores via the trajectories towards 10h cells. To define gene sets for that and to investigate the effects of causal genes with varying strengths of correlation to the overall expression profile on the outcome of Heterogeneity-seq, we chose gene sets with varying correlation strengths to all other genes in the data set. To determine these genes, we first calculated the correlation coefficients for all pairs of genes among all genes with average read counts > 0.5 in 2000 most variable 0h cells. We then ranked genes by the average of their absolute correlation coefficients and binned them in 10 groups according to this order. Group 1 therefore contained genes with weakest correlation to all other genes, whereas group 10 consisted of genes with strong correlation structure.

Next, we calculated sets of trajectories by adding low, average, or high levels of normally distributed noise (mean = 0, standard deviation = 5, 10, or 15) to the distance matrices before trajectory calculation, which changed approximately 21.9, 33.8 and 42.4% of the cell-cell connections compared to the ground truth trajectories, respectively.

### Heterogeneity-sequencing

For the application of Heterogeneity-sequencing (Heterogeneity-seq), consider a time course experiment with *t* time points. We first construct an optimal set of *c* trajectories *T* (see Construction of optimal cellular trajectories). *T* establishes one-to-one relation between cells from the first time point to cells from the last time point.

Next, we calculate a response score (e.g., a module score for a set of genes of interest or viral load) for each cell in the last time point. We then group all these cells into three categories—low, middle, and high responders—based on the value of the response score. The default thresholds for these groups are defined by the quantiles of the response score: low (0–25%), middle (25–75%), and high (75–100%). We assign these response group classifications also to their respective predecessor cells in the first time point.

Now, we apply one of three Heterogeneity-seq approaches to all cells in the first time point using their total RNA expression profile and the assigned response group: (A) testing for differentially expressed genes between the low and high response groups, (B) classification low vs high using a support vector machine (SVM), and (C) predicting response groups based on causal inference (double machine learning). The code was implemented in R (version 4.1.2), and the data set was processed using the Seurat package (version 5.1.0).

In (A), we calculate total RNA log_2_ fold changes and test for differentially expressed genes between low and high responders using the Wilcoxon test. We then apply the Benjamini-Hochberg correction to adjust the p-values.

In (B), our goal is to predict response groups using the expression of a single gene at a time. A gene is a potential pathway modulator if its expression profile alone can be used to accurately predict cellular outcomes. To this end, we use the SVM implementation of the e1071 package (version 1.7-14) with a radial kernel. The classification is encoded as binary, with high responders as the positive class and low responders as the negative class. For each gene, we run 10-fold cross-validation on the whole dataset, obtaining a prediction for each cell without overfitting. The Heterogeneity-seq score for a gene in this variant is the area under the ROC curve (AUC) computed using the pROC package (version 1.18.5). To incorporate additional features for classification (e.g., cell cycle information, viral load, other genes), we can specify a set of informative features *S* and use *g*∪*S* as input features to run the cross-validation and subsequently calculate the increase in the AUC compared to running it with only *S*. We also calculate total RNA log_2_ fold changes for all genes between low and high responders to gauge positive or negative pathway modulation.

In (C), our approach is similar to (B) but a standard SVM is not designed to discriminate between causal, confounding and correlating factors. To take these factors into consideration, we use a causal inference model based on double machine learning with the DoubleML package (version 1.0.1). For each gene *g*, we create a DoubleMLData object using the gene expression data for all genes (including informative features, if applicable), with *g* as the treatment variable and the binarized response variable as the outcome. The DoubleML model is then configured with the DoubleMLData object, 10-fold cross-validation, and two learners ml_g and ml_m from the mlr3 package (version 0.20.2), where both ml_g=“regr.ranger” and ml_m=“regr.ranger”. The model is fitted and for each gene *g* the causal estimate and p-value are extracted. The p-values were adjusted via Benjamini-Hochberg correction.

For all Heterogeneity-seq runs in this study, we only considered genes with a mean expression >= 0.5 in the cells of the first time point. The results were plotted using the ggplot2 package (version 3.5.1).

## Availability of Data and Materials

Raw sequencing data generated for the MCMV infection study has been deposited at GEO under the accession number GSE281259. Raw data from the Dexamethasone time course ^15^ was downloaded from SRA (Accession id SRX5848248)

R notebooks to reproduce all figures (alongside processed data for the Dexamethasone time course ^15^and MCMV infection) are available on zenodo (https://doi.org/10.5281/zenodo.17572588).

The release version, source code and documentation of the HetSeq R package are available on github (https://github.com/erhard-lab/HetSeq). GRAND-SLAM 3.0 is available on github as part of our GEDI framework (https://github.com/erhard-lab/gedi).

## Author information

### Contributions

KB and FE performed analyses, generated figures, and wrote the manuscript, LS and TR performed analyses, KB wrote the source code for the HetSeq package. TH, AWW, KSW, YZ, TK, CT, ML, BKP performed experiments under the supervision of LD and AES, LD and FE developed the conceptual approach of Heterogeneity-seq. All authors read and approved the final manuscript.

## Ethics declarations

Not applicable.

## Ethics approval and consent to participate

Not applicable.

---

**Extended Data Figure 1:**
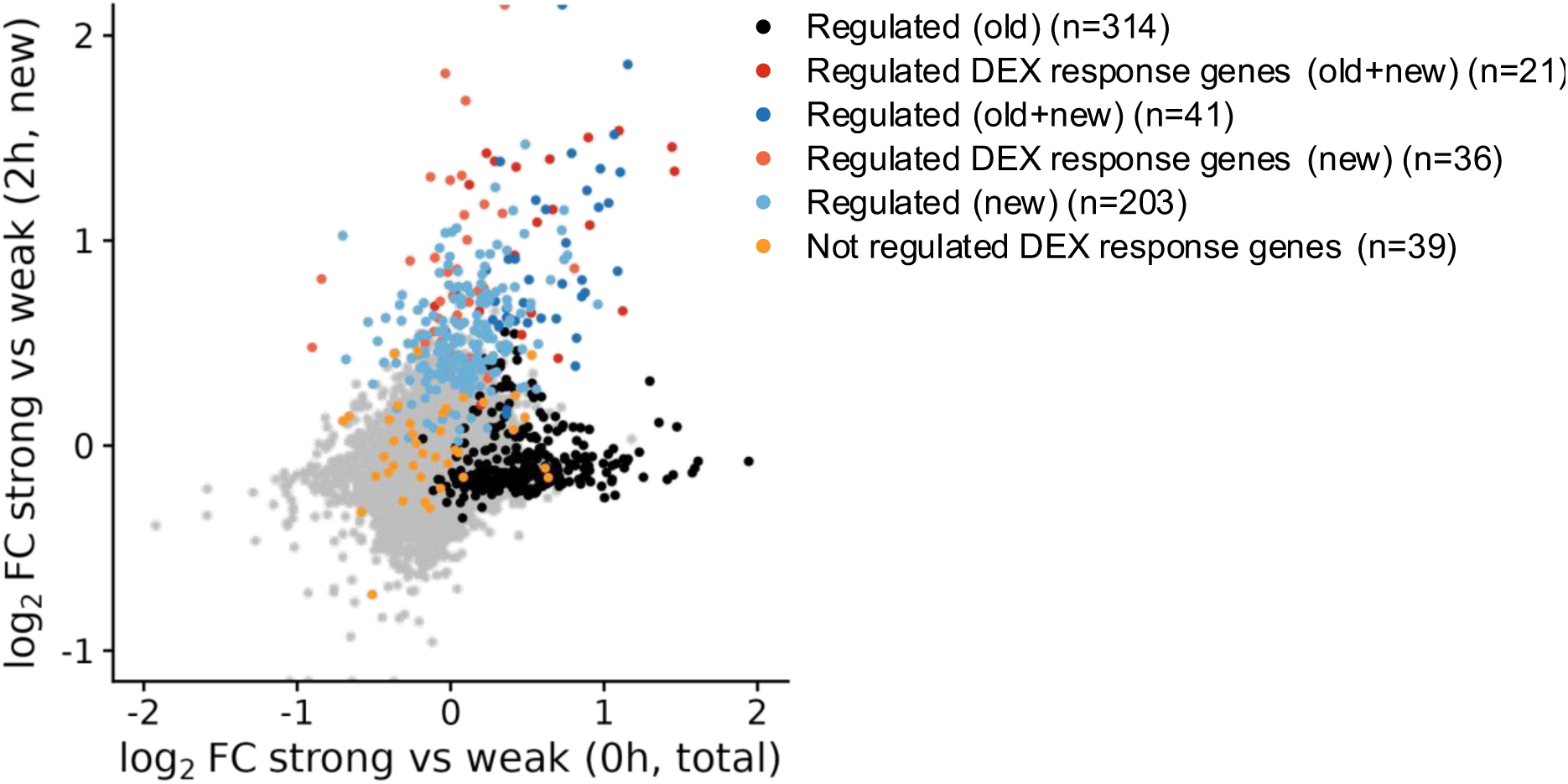
Scatterplot comparing log2 fold changes of pseudobulks of 2h cells (new RNA) versus 0h cells (total RNA) that were connected via trajectories. Similar to Fig. 3e, but regulated genes were not used for constructing trajectories to exclude bias.

**Extended Data Figure 2:**
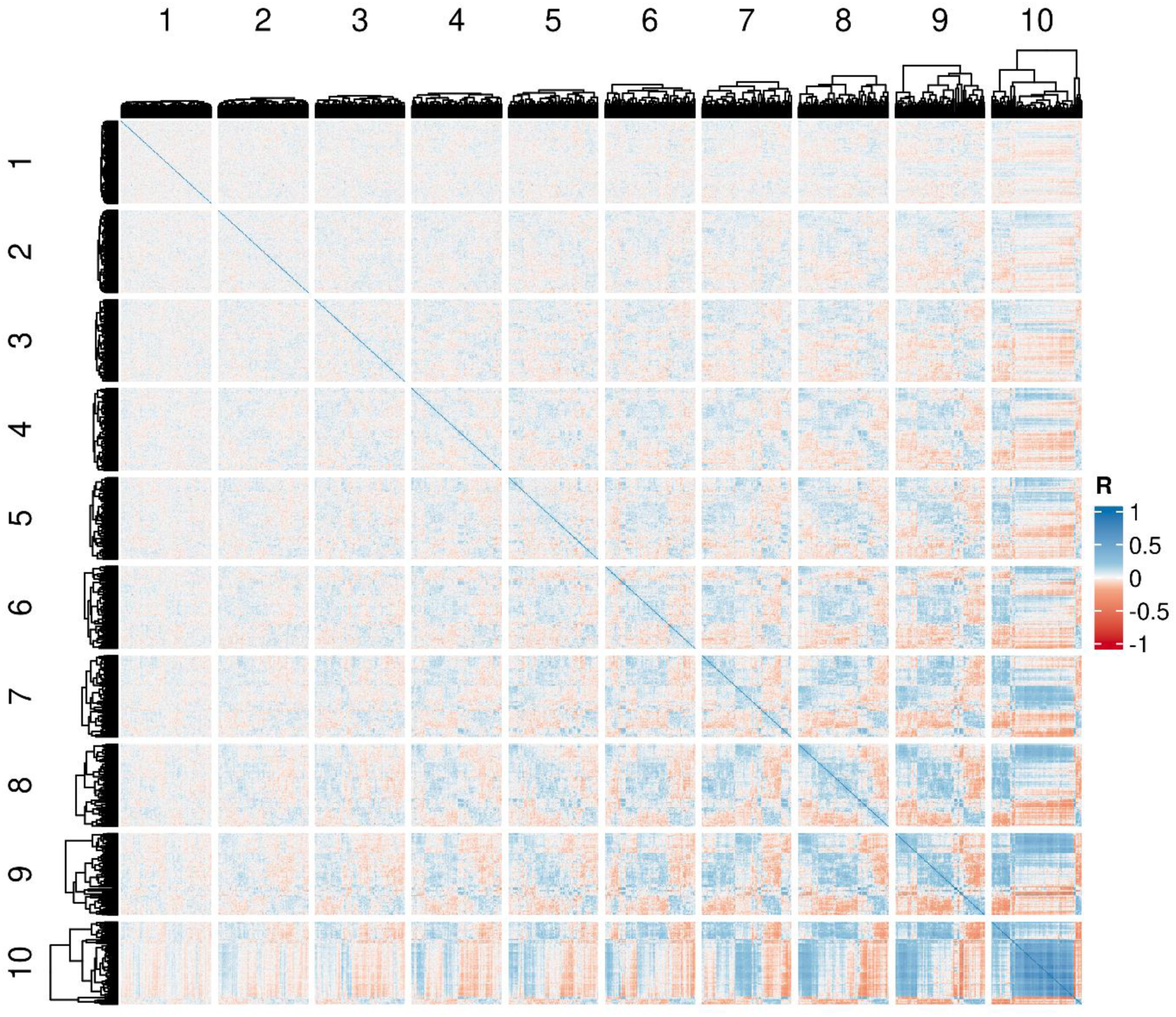
Heatmap showing correlation coefficients (R) for all pairs of genes across 0h cells in the Sci-fate data set. Genes were grouped according to the average correlation coefficient into 10 quantiles as indicated.

**Extended Data Figure 3:**
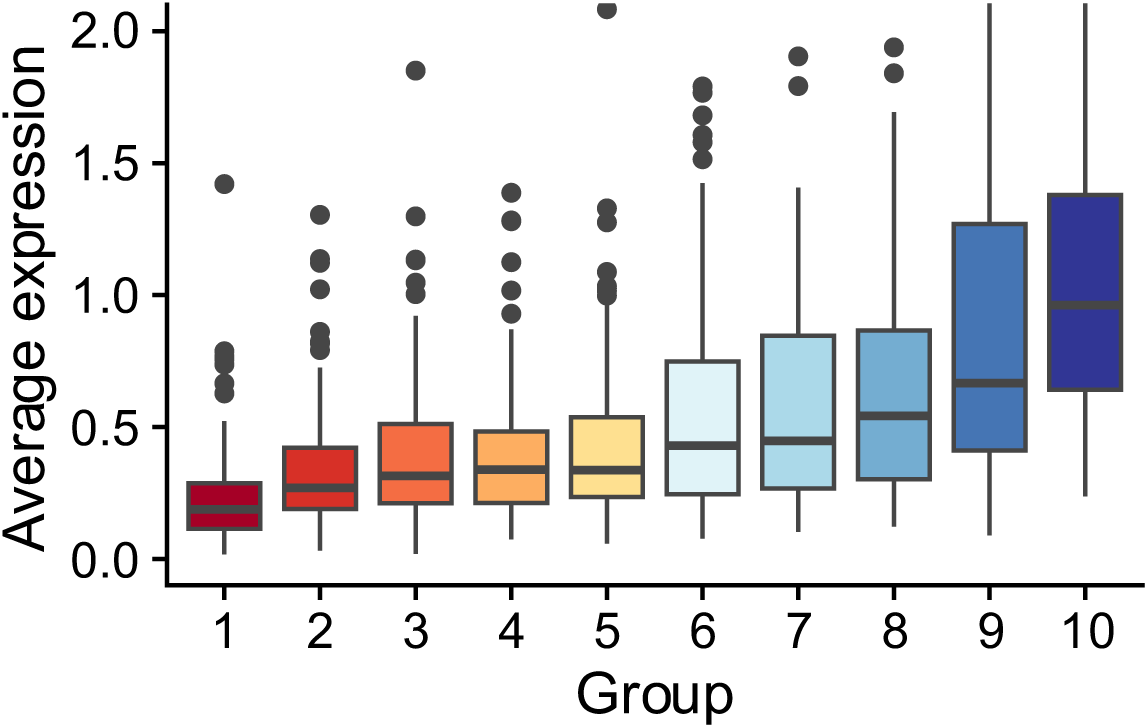
Boxplots (center line, median; box limits, upper and lower quartiles; whiskers, 1.5x interquartile range; points, outliers) showing the distribution across genes of expression values for the 10 groups defined in **Extended Data** Fig. 2. Each value in these distributions is the average expression value computed across all 0h cells (log-normalized) of the Sci-fate data set.

**Extended Data Figure 4:**
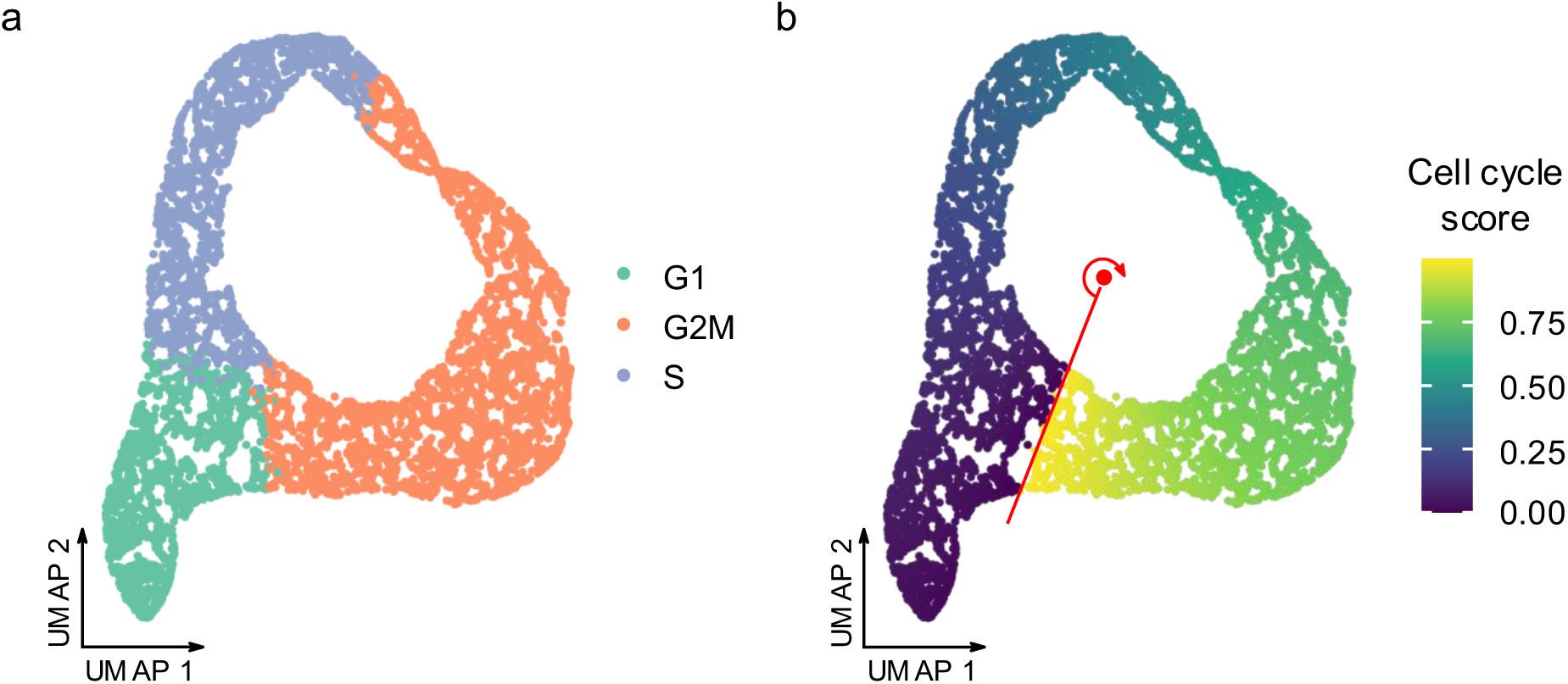
Definition of a positional cell cycle score. **a** UMAP representation of DEX-treatment data set showing Seurat’s classification of cell cycle phases **b** Definition of the cell-cycle score by the cell position around a manually defined cycle center (red point).

**Extended Data Figure 5:**
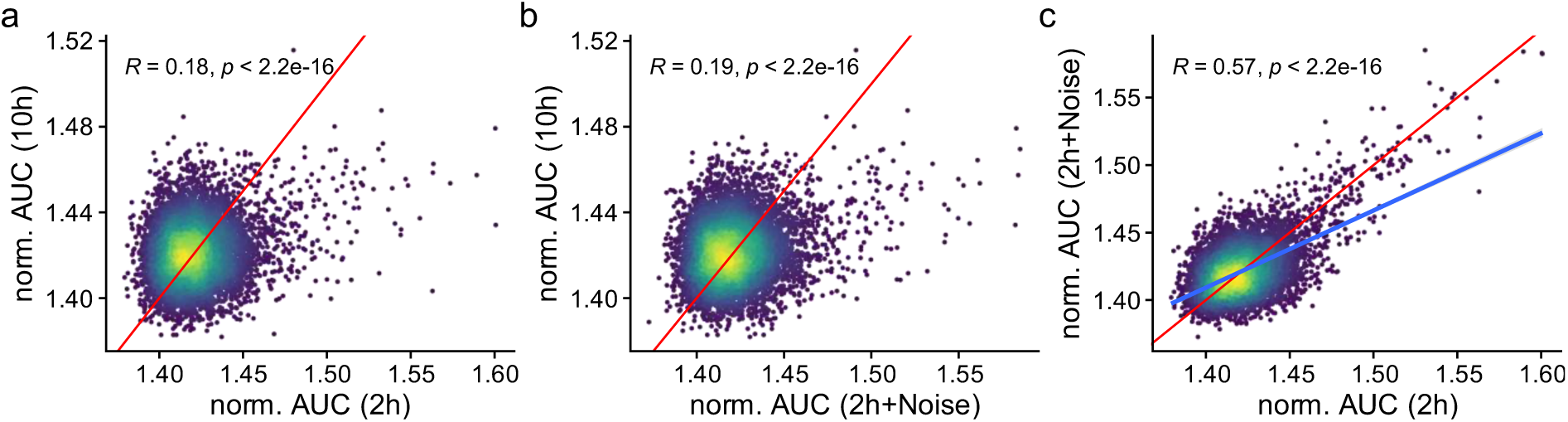
The impact of trajectory noise on Heterogeneity-seq results. Scatter plots comparing the predictive value (area under the receiver operating curve - AUC) for each gene for two different Heterogeneity-seq runs. In all cases, the cell cycle was used as an additional feature, and the AUC values were normalized by dividing by the sum of all AUCs for that run and multiplying by 10,000. The origin line (red), Pearson correlation and p-values (two-sided approximate t-test) are indicated. **a** Comparing 2h versus 10h. For the x-axis, the 2h DEX score was used to define low and high responder cells, whereas for the y-axis, 10h cells were used. **b** Comparing 2h including noise versus 10h. For the x-axis, the 2h DEX score was used to define low and high responder cells, whereas for the y-axis, 10h cells were used. For the 2h Heterogeneity-seq run, normally distributed noise was added to the distance matrix of trajectory inference. **c** Comparing 2h including noise versus 2h without noise. A regression line (blue) is indicated.

**Extended Data Figure 6:**
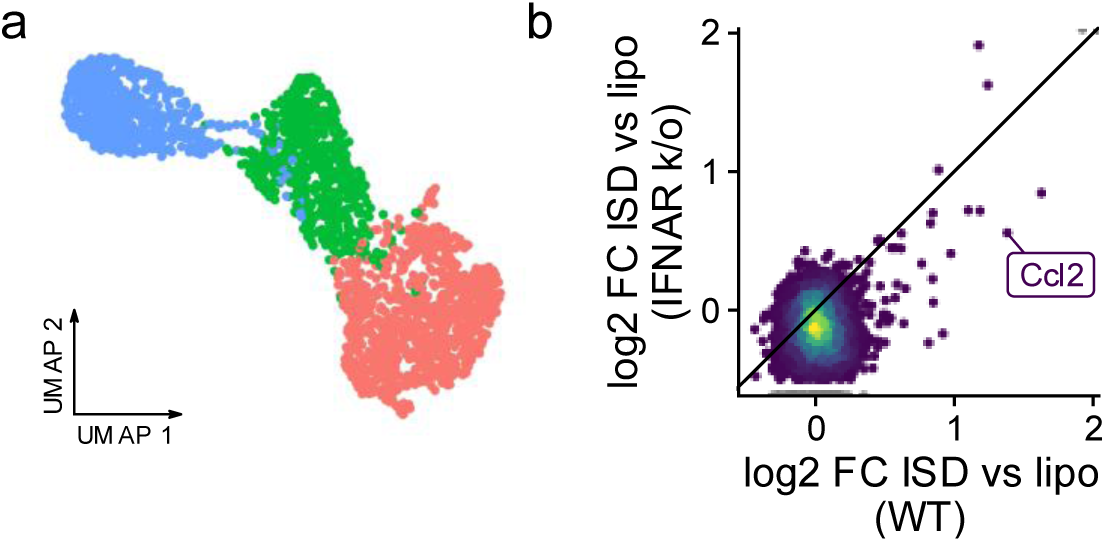
MCMV scSLAM-seq data. **a** UMAP embedding with cells colored according to Louvain-clustering with resolution 0.1 on the first ten principal components. We used a set of n=205 genes at least 2-fold upregulated upon MCMV infection in NIH-3T3 cells (Erhard et al, Nature 2019) for computing the principal components. **b** Scatterplot comparing the log_2_ fold change (log2 FC) of wild-type mouse fibroblasts (WT) against IFNAR knockout (k/o) cells. For both cells the log2 FC was computed from cells treated with immunostimulatory DNA (ISD) versus control cells treated with lipofectamine. The induction of Ccl2 in both conditions is indicated. Data taken from Schwanke (2023), Supplementary Table of GSE224855.

**Extended Data Figure 7:**
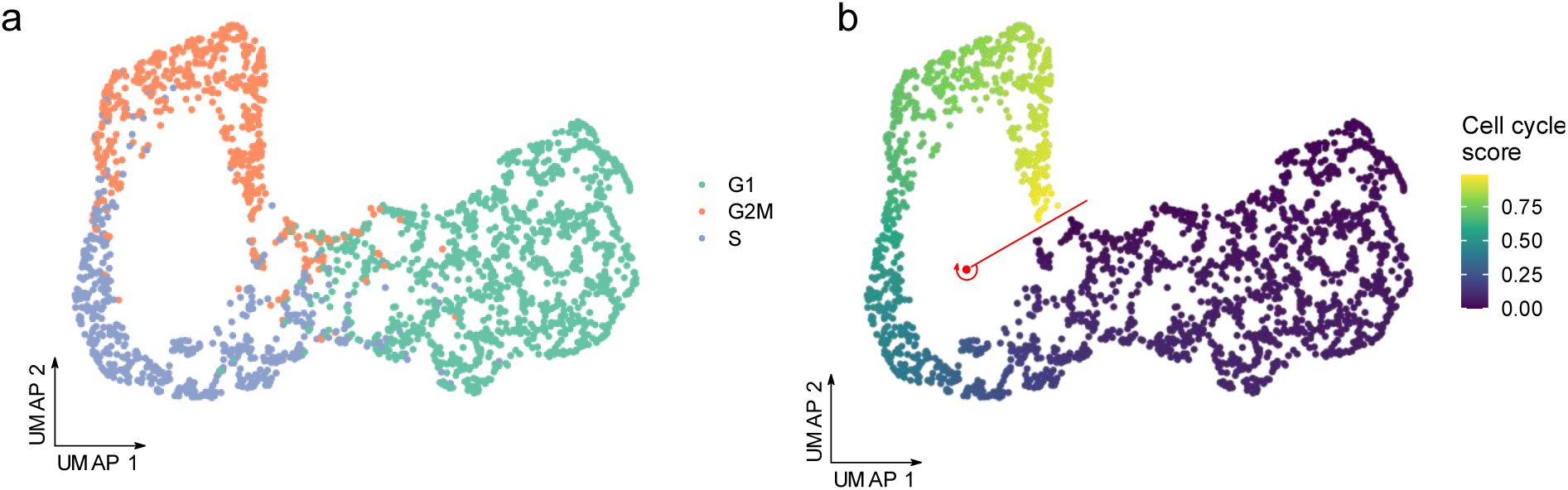
Definition of a positional cell cycle score. **a** UMAP representation of the MCMV data set showing Seurat’s classification of cell cycle phases **b** Definition of the cell-cycle score by the cell position around a manually defined cycle center (red point).

